# Dynamics of microglia-glioblastoma crosstalk at the far infiltration zone

**DOI:** 10.1101/2024.09.23.614016

**Authors:** Felix C. Nebeling, Falko Fuhrmann, Manuel Mittag, Fabrizio Mussachio, Nala Gockel, Lea L. Friker, Sonia Leonardelli, Severin Filser, A Deli, Miriam Stork, Daniele Bano, Torsten Pietsch, Simona Parrinello, Michael Hölzel, Ulrich Herrlinger, Paolo Salomoni, Martin Fuhrmann

**Author notes:** These authors contribute equally (Paolo Salomoni, Martin Fuhrmann). Corresponding Authors: Dr. med. Felix C. Nebeling, Prof. Dr. Paolo Salomoni, Prof. Dr. Martin Fuhrmann.

## Abstract

The interaction of glioblastoma (GB) and microglia is critical due to its implications for tumor progression, immune response modulation, and potential therapeutic strategies. However, the role of microglia in GB pathogenesis remains unclear, especially regarding the *in vivo* dynamics of their interplay. Performing three-photon imaging in an autochthonous, immunocompetent mouse GB model, we examined tumor/microglia dynamics within previously inaccessible regions at the GB far infiltration zone in the *corpus callosum*. Initially, microglia increased tissue surveillance upon encountering GB-cells in sparsely infiltrated areas. In contrast, when GB-cell density increased, microglia reduced surveillance, suggesting a biphasic response to tumor invasion. Additionally, microglia were not uniformly attracted to infiltrating GB-cells; only a subset moved directionally toward them within a defined spatial range. This study provides insight into the heterogeneity of the immune response to tumor invasion and the dynamics of microglia-GB interactions in vivo.

**Graphical abstract:** 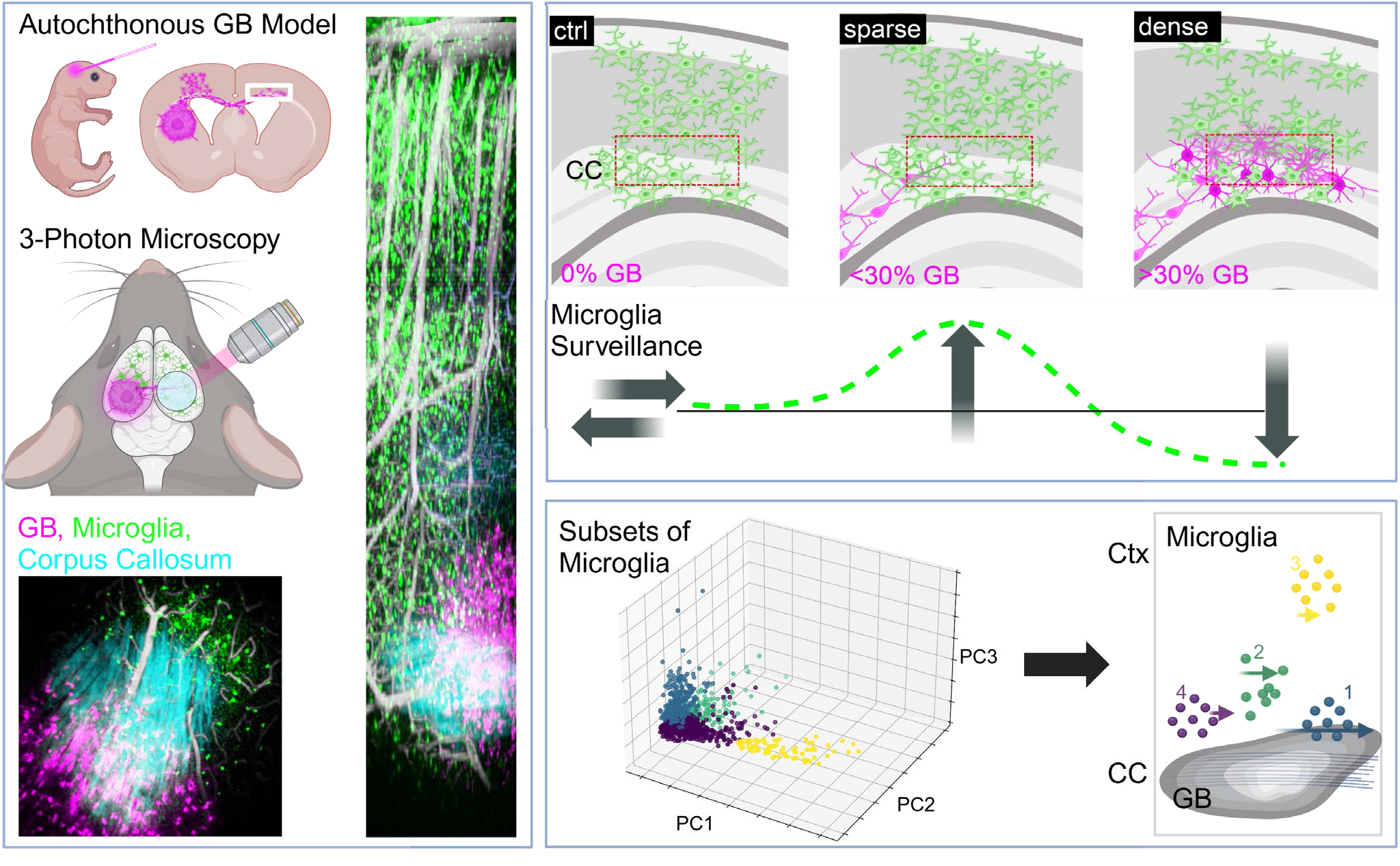

## Introduction

Glioblastomas (GBs) are the most common and most malignant primary brain tumors in adults.^1^ Despite extensive therapeutic intervention, including surgery, radiotherapy, and chemotherapy, the median survival for patients in trial cohorts remains limited to about 17 months.^2^ GBs typically form a hypoxic bulky tumor with a surrounding infiltration zone. The diffuse infiltration of GB-cells into the adjacent parenchyma is a hallmark of GB pathology and can be spatially subdivided into a near and far infiltration zone. Recently, the near infiltration zone has been described in mouse models of GB as the area where GB-cells reside within 0,5 – 2 mm distal to the tumor bulk.^3,4^ The far infiltration zone can be defined as the dynamically changing – in space and time – invasion front of GB-cells more than 2 mm apart from the tumor bulk. Increasing evidence suggests that GB-cells in these tumor niches are functionally distinct.^5–9^ In addition to differences in intrinsic features of GB-cells depending on tumor location, the tumor microenvironment (TME) has also been suggested to be distinct within the different niches,.^10,11^ GBs develop within a complex tissue microenvironment composed of diverse cell populations including infiltrating and resident immune cells, which may comprise up to 30-50% of the non-neoplastic cells within the TME.^12,13^ Tumor-associated macrophages and microglia have been suggested to play a tumor-supportive role in glioma pathogenesis, as they are prone to release a variety of factors facilitating tumor growth and surrounding tissue necrosis by establishing an immunosuppressed microenvironment.^14–17^ However, we lack a comprehensive understanding of microglia dynamics in GB, in particular of whether they present morphological and functional heterogeneity in relation to distinct tumor niches. Under physiological conditions, microglia constantly surveil their microenvironment with highly plastic fine processes, a phenomenon termed microglial motility.^18,19^ This surveillance capacity enables microglia to quickly detect and respond to even subtle changes within their microenvironment. Although key drivers of microglial motility have been identified under homeostatic conditions and certain brain pathologies, our understanding about microglial dynamics during GB-cell infiltration is limited. Most studies on microglia in this context rely on static histology or *in vitro* models and focus on the tumor bulk.^10,14,20^ Despite the critical role played by infiltrating GB-cells in the relapse upon surgical resection,^21,22^ the behavior and features of GB-cells at the far infiltration zone of GBs remain insufficiently explored. Therefore, there is an urgent need for *in vivo* assays of the far infiltration zone, particularly concerning microglial dynamics, which have not been investigated in this niche *in vivo* so far.^23–25^ However, studying microglial dynamics in these brain areas remains challenging, because of (i) anatomical inaccessibility for intravital two-photon microscopy and (ii) paucity of faithful GB mouse models. The shortcomings of most commonly used GB mouse models, GL261 and patient-derived xenografts (PDXs), are excessive immunogenicity, lack of diffuse infiltration patterns^26^ and incomplete immune system in the case of PDX models.^27^ These limitations may explain why many immunotherapeutic drugs successfully tested in preclinical settings later failed in clinical trials.^28–32^ Therefore, we need more representative mouse models that recapitulate both, the diffuse infiltration and immunogenicity to foster informed GB immunotherapies. Accordingly, we and others have developed autochthonous models of GB based on *in vivo* electroporation, which bear promise for investigation of GB/TME dynamics and identification of new tumor vulnerabilities. ^3,33–36^ Regarding anatomical accessibility for intravital microscopy, one of the major routes of GB-cell infiltration is the *corpus callosum* (CC).^21,37–40^ The CC in adult mice is deeply embedded under the cortex, approximately 800 µm below the pial surface. Conventional two-photon (2P) microscopy does not enable non-invasive imaging of this anatomical structure, thus precluding investigations of the far infiltration zone.^41^ Recently, three-photon (3P) microscopy has emerged as a promising tool for deep brain intravital imaging, allowing for non-invasive visualization of cellular structures as deep as 1100-1600 µm below the brain surface, depending on the brain area.^42,43^ In a recent study, 3P microscopy has been applied to investigate GB-cells inside the CC, however in an immunodeficient PDX model.^44^ Thus, GB-cell and microglial dynamics at the far infiltration zone within an immunocompetent TME remain unknown.

Here, we apply longitudinal 3P-imaging to a somatic model of IDH^WT^ Mesenchymal (Mes) GB.^3^ Our *in vivo* work reveals that in the CC distinct morphological GB features correlate with invasion velocity. Furthermore, we show differential regulation of microglial motility and directionality of movement in response to increasing GB tissue burden. Overall, this study provides understanding of the *in vivo* heterogenous response of innate immune cells to GB-cell infiltration, which might also be relevant for other brain pathological states displaying alterations of microglia function.

## Results

### GB-cell velocity inversely correlates with number of tumor microtubes at the far infiltration zone

In order to study diffuse infiltration patterns within an intact TME, we chose an autochthonous GB model based on *in vivo* electroporation, which we and other have previously described **(Figure 1)**.^3,33–36^ We injected a DNA mix containing plasmids carrying a piggyBac-integrable fluorescence reporter, episomal Cas9 and specific gRNAs for triple inactivation of the *Nf1*, *Trp53* and *Pten* tumor suppressors, the main driver of Mes GBs, ^45^ into the ventricular space at postnatal day two. Upon electroporation, fully penetrant Mes GBs developed in the forebrain, as previously reported.^3,33,36^ With a latency of 2-3 months post injection (p.i.), mice developed high-grade, diffusely infiltrating gliomas **(Figure 1A–D, Video S1)**. GB-cells reliably infiltrated the contralateral hemisphere via the *corpus callosum* constituting the far infiltration zone **(Figure 1B-D)**. Bulky lesions were enriched in CD44 as well as in cells of myeloid lineage expressing CD11c recapitulating a key histological feature of human Mes GB, ^46–49^ which our model replicates **(Figure 1E)**. We further found evidence for ectopic proliferation with the tumor bulk (28,5% Ki67 positive nuclei vs. 2.25% within the healthy tissue; **Figure1F)**. Together, these findings confirm the ability of this electroporation-model to recapitulate invasive Mes GBs, offering the opportunity to investigate tumor spreading in living animals. As intravital microscopy methods based on 2P-imaging do not allow to visualize infiltration of GB-cells within deeper brain areas relevant for invasion, such as the CC, we performed 3P-imaging, which provides access more than a millimeter deep in the brain.^42,43^ We implanted cranial windows 6 weeks p.i. over the somatosensory cortex of the contralateral hemisphere relative to the side of tumor induction, as visualized by the fluorescent tdTomato signal from beneath the skull **(Figure 2A-C)**. The CC is a deeply embedded anatomical structure, which in mice resides approx. 800 µm below the brain surface. After a 4-week recovery phase, intravital microscopy was conducted for up to 4 hours **(Figure 2A)**. Imaging the same cortical vasculature in wildtype mice following i.v. Texas Red^TM^ Dextran injection confirmed the superiority of 3P-imaging, especially in terms of penetration depth as compared to 2P-imaging **(Figure 2D)**. In our set-up, 2P-excitation at λEx=920nm was better suited for imaging the mouse cortex up to approx. 400 µm below the pial surface. Below that, 3P-excitation at λEx=1300nm was superior, enabling to image deeper than one millimeter **(Figure 2D and 2E)**. To visualize the deeply embedded CC, we recorded the third-harmonic generation (THG) signal with excitation at λEx=1650nm **(Figure 2F and 2G, Video S2)**. THG occurs at material or tissue interfaces with different optical properties such as local changes in refractive index or nonlinear susceptibility. The CC primarily consists of myelinated axonal fiber tracts,^50^ which are rich in lipids and produce a strong THG signal if subjected to 3P-excitation.^51^ It was reliably possible to excite fluorescence (λEx=1650nm) of infiltrating tdTomato^+^ GB-cells inside the CC of adult mice **(Figure 2G)**. Next, we measured the migration and morphological properties of individual GB-cells in the CC over several hours **(Figure 2H and 2I)**. Close to the tumor bulk GB-cells have been shown to exhibit different migration velocity in relation to their tumor microtubes.^9^ Tumor microtubes (TMs) are small protrusions of GB-cells that are important for physiological interconnection and communication between tumor cells and with the surrounding TME.^52^ Accordingly, we found that GB-cells with few TMS migrated longer and with higher velocity at the far infiltration zone in our autochtonous model. GB-cells with many TMs traveled shorter distances and were slower **(Figure 2I-L, Video S3)**. If GB-cells were divided into cells with ≤ 4 TMs and those with > 4 TMs, we observed a significantly longer traveled distance for GB-cells with fewer than 4 TMs **(Figure 2J** and **2K)**. GB-cells with less than 4 TMs migrated with a velocity of 15.3 µm/h, which was 72% faster than GB-cells with more than 4 TMs (8.9 µm x h^-1^; **Figure 2L**). Moreover, we identified an inverse correlation between the number of TMs and the distance traveled by GB-cells (r = -0.47; **Figure 2M**). Overall, these findings provide novel insights into the *in vivo* behavior of GB cells, suggesting a link between number of TMs and GB-cells’ traveled distance and velocity at the far infiltration zone.

**Figure 1.**
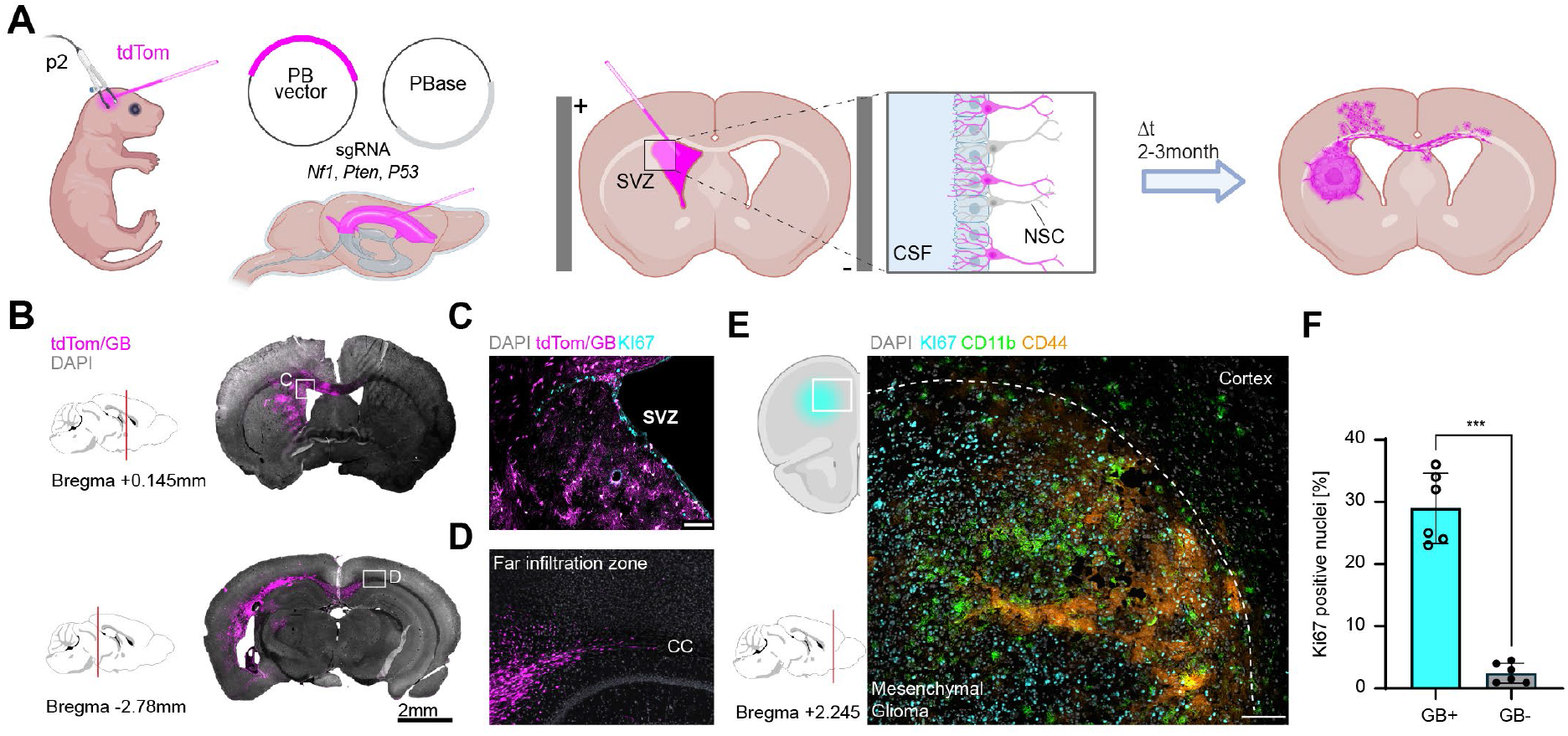
CRISPR/Cas-mediated autochtonous model of adult mesenchymal Glioblastoma. **A** Schematic of the CRISPR/Cas-mediated generation of an autochtonous Glioblastoma (GB) model with mesenchymal signature (sgRNAs for Nf1, Pten, P53). A PB vector containing three gRNAs alongside the tdTomato fluorescent reporter and a second plasmid with the Cas9 and PiggyBase (PBase) under control of the hGFAPmin promoter is injected into the lateral ventricle of P2 pups. After electroporation mice develop highly aggressive GBs after a latency of 2-3 month with high penetrance. **B** Exemplary images showing the diffuse infiltration of tdTom+ GB-cells in an adult mouse (2 months). Note the long ranging infiltration via the *corpus callosum* crossing to the far infiltration zone of the contralateral hemisphere. **C** Close up image of the targeted subventricular zone (SVZ). **D** Close up image of GB-cells infiltration to the contralateral hemisphere via the CC constituting the far infiltration zone. **E** Excerpt of the tumor bulk and the near peritumoral tissue highlighting the immunoreactivity for CD44 and CD11c resembling the mesenchymal subtype of human GB. **F** Quantification of Ki67 positive nuclei inside (GB+) and outside of Glioma bearing tissue (GB-). Paired t-test, two tailed; *p*=0.0002; *t*=10.00; *df*=5; n=6 samples from n=3 mice. \*\**p*<0.01; Scale bars 2mm (B, left panel; C left panel), 100μm (B, left; D) 50μm (C, right panel)

**Figure 2.**
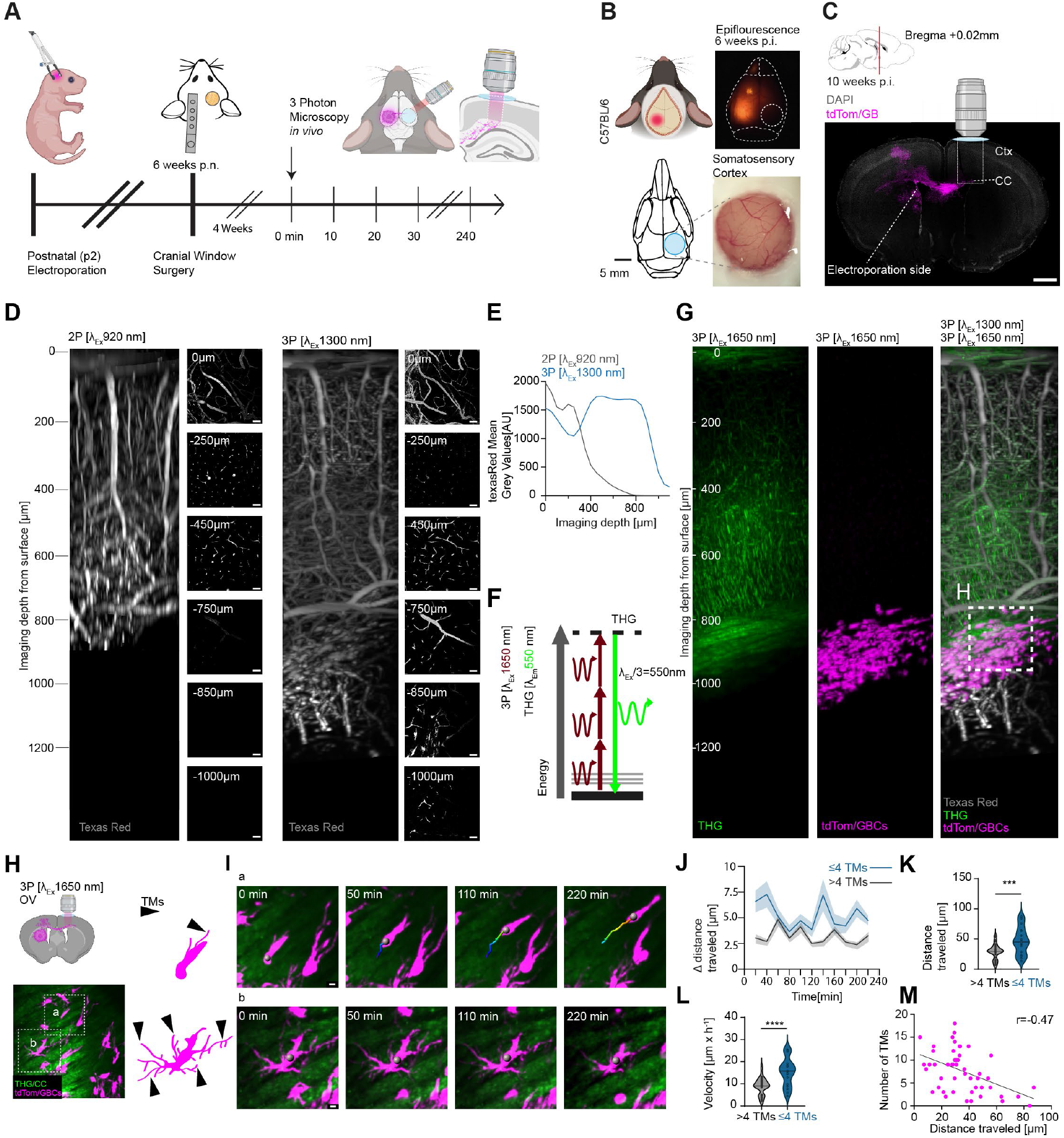
Number of GB-cell protrusions are predictive of invasion velocity via *corpus callosum*. **A** Schematic timeline of the experimental procedure. At P2 mice are injected with the CRISPR/Cas+Transposom plasmid mix and electroporated. After 6 weeks a cranial window is being implanted. To allow the mouse to recover and let acute inflammation subside, a 4 week-period is given before the onset of intravital microscopy. During the initial imaging paradigms consecutive time-lapse imaging was performed with a 10 minute-interval for about 4 hours. **B** The bulky tumor is located using epifluorescence during surgery. Subsequently, the craniotomy is performed over the contralateral hemisphere, where no tumor is visible yet. **C** This allows *in vivo* imaging of the far infiltration zonein the *corpus callosum* and adjacent cortical tissue **D** 3D reconstruction of the same volume with Texas Red (70-kDa dextran-conjugated fluorescent dye) labeled vasculature imaged with 2P (λ_Ex_=920nm) and 3P (λ_Ex_=1300nm) excitation. The fluorescent intensity of individual layers was normalized for better visualization. Imaging of vasculature was sufficient until ca. 450 µm in 2P mode and up to 1000 µm using 3P imaging. **E** Quantification of mean Texas Red intensity at increasing imaging depths. 2P excitation is superior over 3P within the first 400 µm below the pial surface. At larger depths 3P microscopy yields much more signal. **F** Third harmonic generation (THG) is generated at structural interfaces with high nonlinear susceptibility. Here the energy of one excitation photon is combined to a single photon with 1/3 of the excitation wavelength. When excited at λ_Ex_=1650nm the THG signal will have 1/3 of the wavelength which can be detected in the green-yellow fluorescent spectrum (λ_Em_=550nm). **G** Vessels and white matter tracks are visualized label-free via Third Harmonic Generation using an excitation wavelength of 1650 nm. The tdTomato+ GB-cells are prominently visible deep within the CC. Combing 1300 and 1650 nm excitation wavelengths within the same brain volume, one can co-visualize the THG, tdTom and Texas Red. **H** Imaging of the far infiltration zone in the CC of the contralateral hemisphere (in respect to the tumor bulk). Cells display great heterogeneity in terms of cellular protrusions with none or few putative Tumor microtubes (TMs) to a multitude of cellular protrusions, marked in schematic with black arrows. Notice, these cellular states coexist directly next to each other. **I** Exemplary time-lapse images of a GB-cell with a rod-like shape (upper panel) and little number of TMs. Lower panel: GB cell with a multitude of cellular protrusions. Definition of TM: ≥10 µm length and 0.5-3 µm diameter. The somata are marked with a sphere and the movement track of the cell depicted. **J** GB-cells traveled distances plotted over individual time points, >4TMs grey, ≤4 TMs blue. **K** Comparison of traveled distance between GB-cells with >4 or ≤4 TMs; GB-cells with ≤ 4 TMs travel longer distances in the same time then cell with >4TMs (29.09 µm vs. 47.02 µm ±5.03 µm) Unpaired *t*-test, p=0.0008 t=3.566; *df*=47 **L** Average velocity of GB-cells with >4TMs or ≤4 TMs during 1h (8.93 µm vs 15.26 µm ±1.46 µm); unpaired *t*-test, two tailed*, p*<0.0001, df=4.341, *df*=47 **M** Correlation of traveled distance of GB-cells during a 3h imaging period and their number of TMs; n=49 cells; Pearsons *r*= -0.47, *R^2^*=0.22, p=0.0006. ***p<0.001, ****p<0.0001; Scalebars: 500 µm (C), 50 µm (D), 10 µm (I)

### GB-cell infiltration velocity depends on anatomical location

After measuring migratory properties of GB-cells within the CC over hours, we investigated GB-cell migration at the far infiltration zone over several days. Intravital microscopy was carried out repeatedly in 2.5 - 3 months old GB mice for three to four consecutive days with 24-hour intervals **(Figure 3A)**. To monitor migration of GB-cells within the CC and in the adjacent cortex, we recorded large z-stacks spanning approximately one millimeter, which covered the entire cortical column and the CC **(Figure 3B)**. Imaging the same volume repetitively revealed an increase in GB-cell density, nearly doubling every 24h; 97% increase from 0-24h and 66% from 24-48h (n0h=585 ±171 GB-cells per volume, n24h=1154 ± 357 GB-cells and n48h=1923 ±634 GB-cells/mm^3^) **(Figure 3, right panel insert, Video S4)**. We found that GBCs residing in the cortex were slower and migrated shorter distances in 24 hours than in the CC **(Figure 3C-E)**. The mean velocity of GB-cells in the CC was significantly increased and about two times higher than in the cortex **(Figure 3E**). The relative frequency distribution of distances traveled per GB-cell revealed that there might exist two subsets of GB-cells, a large fraction with a velocity of up to 20 µm/d and a smaller fraction with higher velocity of up to 100 µm/d **(Figure 3F)**. 87.5 ±4.5 % of GB-cells in the cortex were slow (< 20 µm/d) compared to 42.8 ±11.2 % in the CC. Whereas 57.2 ±11.2% of GB-cells in the CC were fast (≧ 20 µm/d) compared to 12.5 ±4.5 % in the cortex **(Figure 3G and 3H)**. Overall, these data reveal: i) higher migration velocity of GB-cells within the CC compared to the cortex; ii) heterogeneity of GB-cell fractions (slow/fast) within these two brain regions.

**Figure 3.**
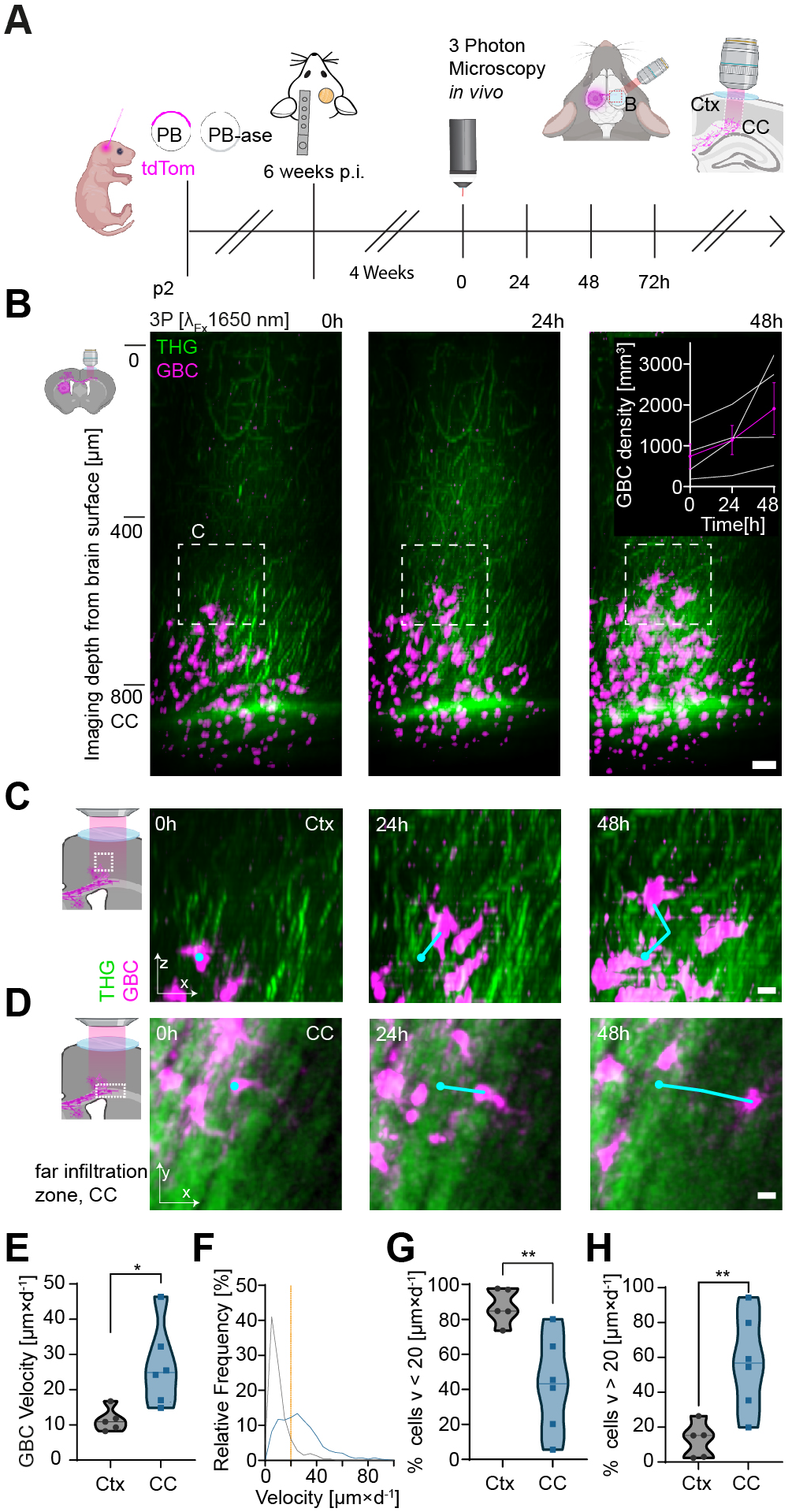
GB-cells show higher velocity in the CC compared to Cortex. **A** Schematic timeline of the experimental procedure as described before. At P2 mice are injected with the CRISPR/Cas+Transposom plasmid mix and electroporated. After 6 weeks a cranial window is being implanted. To allow the mouse to recover and let acute inflammation subside, a 4 week-period is given before the start of intravital microscopy. Imaging was performed with a 24h-interval for 3-4days. **B** Side view of z-volume on of the contralateral hemisphere in respect to the tumor bulk. Images were acquired at λ =1650nm spanning approx. 1000 µm from the pial surface through the CC. Within 3 consecutive days we saw an increase of GB-cell density with GB-cell numbers nearly doubling per 24hs. **C, D** Exemplary images of GB-cells migrating inside the Cortex (C) and CC (D) at the far infiltration zone. **E** Comparison of velocity between GB-cells in cortex and CC. (n=5 mice Ctx and n=6 mice CC; unpaired *t*-test, two-tailed, p=0.0188). **F** Relative frequency distribution of cortical and CC-associated GB-cells in respect to their 24h velocity. Cell movement was binned to 5 µm units; n=614 GB-cells in cortex and n=538 CC-associated GB-cells from n=5 mice. **G** Discrimination of cells with an average velocity of < 20 µm x d^-1 1^ in Ctx and CC (unpaired *t*-test, two-tailed, *p*=0.0075, n=5 mice). **H** Discrimination of cells with an average velocity of > 20 µm x d^-1^ in Ctx and CC (unpaired *t*-test, two-tailed, *p*=0.0075, n= mice). Scale bars 50 µm (B), 20 µm (C,D) * p<0.05, ** p<0.01

### Microglial surveillance displays a biphasic response to GB-cell infiltration

Changes in microglia function and state have been implicated in GB pathogenesis,^14,15,17^ with some studies reporting an association between increased microglia activation markers and poor prognosis in human patients.^53,54^ In agreement with this, we found a general increase of CD68^+^ cells within tumor infiltrated human tissue **(Figure 4A and 4B)**. Specifically, in GB-cell infiltrated areas, CD68^+^ cells made up 38.4 ±4.5% compared to 11.4 ±1.3% in GB-cell negative control areas. Interactions of microglia with GB-cells have mostly been studied with static histological methods. We wanted to analyze the dynamics of microglia/GB-cell interaction, in particular at the far infiltration zone. To this end, we employed *in vivo* electroporation of *Cx3cr1^GFP^*/^wt^ transgenic mice that express green fluorescent protein (eGFP) in microglia,^55^ followed by intravital 3P-imaging **(Figure 4C)**. We simultaneously recorded eGFP-expressing microglia, tdTomato-positive GB-cells and myelinated CC fibers in a cuboid volume spanning the entire cortex and CC **(Figure 4D, Video S5 and S6)**. First, we measured morphologic parameters of microglia within the CC in the absence (GB^–^) and presence (GB^+^) of GB-cells **(Figure 4E)**. By performing Sholl-analysis of microglia ramification, we revealed reduced intersections of microglial branches with radiuses around the cell soma. The number of intersections were reduced in relation to increasing distance from the microglial soma in GB^+^ compared with GB^–^ samples, indicating shorter and less branched processes of microglia and an amoeboid cell shape **(Figure 4F and 4G)**. After determining static morphological differences, we measured the dynamics of microglial fine processes during GB progression. We compared three different stages of GB-cell infiltration within the CC: ctrl, sparse and dense (0%, <30% and >30% GB-cell coverage; **Figure 4H**, upper panel). We located the CC via the THG signal generated at λEx=1650nm (**Figure 4H**, second panel). Microglial motility was measured in a repetitive manner acquiring z-stacks every 5 minutes spanning 150 µm through CC **(Figure 4H, S7-10)**. Microglial process turnover rate (TOR), gained, lost and stable fraction were calculated as previously described.^56,57^ The number of all gained (green) and lost (red) pixels divided by the overall number of visible pixels, in which non changed pixels would appear yellow, is the turnover rate (TOR) of microglial fine processes **(Figure 4I)**. In the absence of GB-cells within the CC, microglial processes displayed an average TOR of 40 ±1.2%. Upon sparse infiltration of the CC microglia significantly upregulated their surveillance by 17% (47.1

**Figure 4.**
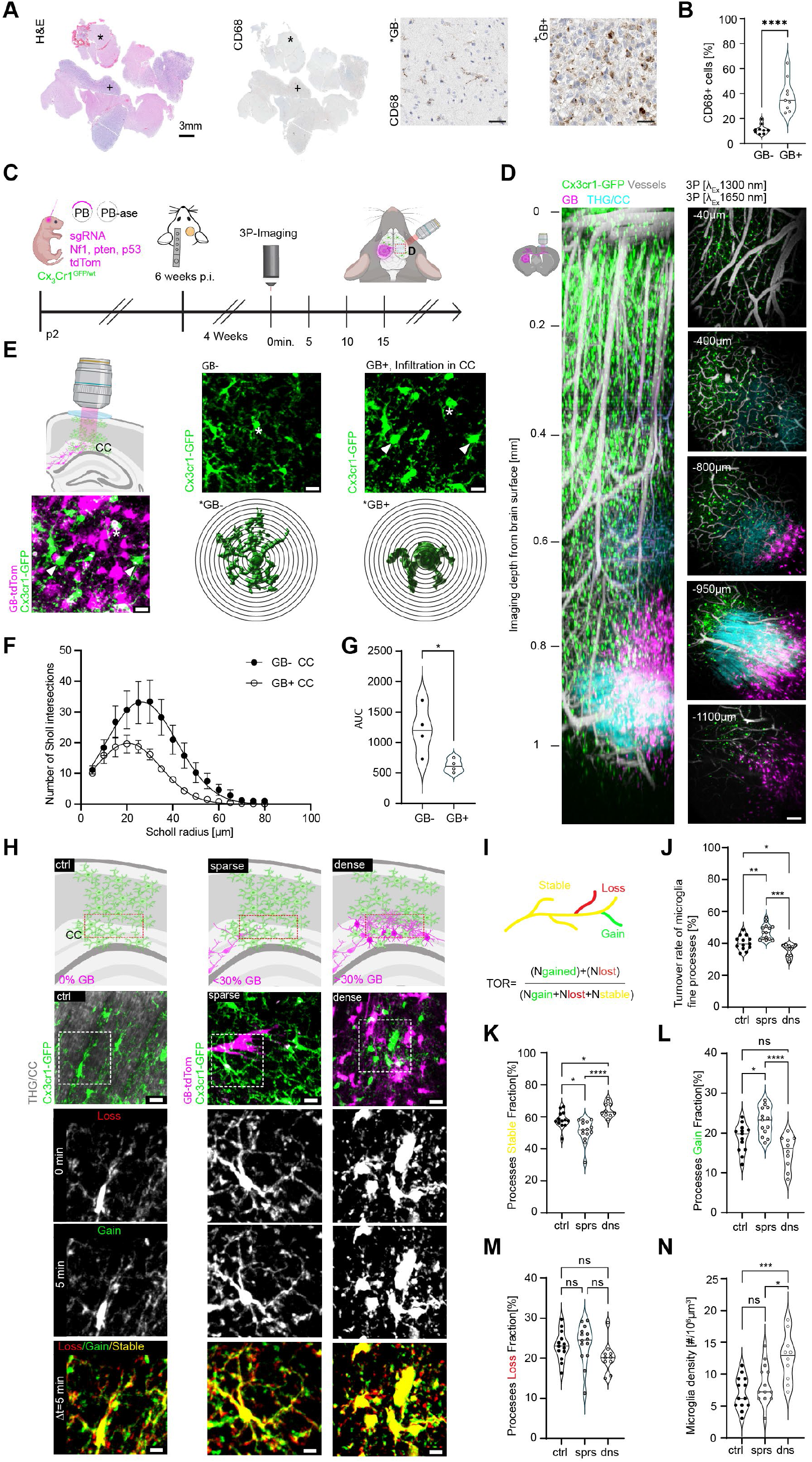
Differential regulation of microglial surveillance upon increasing levels of glioma infiltration in *corpus callosum*. **A** Overview of human Glioblastoma FFPE consecutive resection samples stained H&E and against KP1 (CD68), showcasing tumor dense and non-infiltrated zones as indicated by asterisk (non-infiltrated) and a plus (+, GB infiltrated tissue). The right panel shows the respective zoom images of non-infiltrated and infiltrated areas highlighting the changes of CD68 cell numbers and putatively their shape upon GB infiltration in patient resected samples. **B** Quantification of CD68 positive cells in GB free versus infiltrated tissue, n=9 samples with a total of n=170258 cells, GB-11.42± 1.28 µm vs GB+ 38.38±4.46 µm; paired t-test, two-tailed, p<0.0001, t=7,235, df=8. **C** As a recipient we use the Cx_3_Cr-1 ^GFP/wt^ mouse line. At P2 the CRISPR/Cas plasmid mix (sgRNAs against Nf1, pten and p53 and a Transposom system for fluorescent visualization) is injected into the lateral ventricle with consecutive electroporation and cranial window surgery as shown previously (Fig.1,2). About 3 months after injections intravital 3P microscopy is performed. **D** 3D reconstruction of a z-stack spanning approx. 1150 µm through the entire cortex and the deep-seated corpus callosum on the contralateral side in respect to the tumor bulk/the electroporation side. Two subsequent z-stacks were acquired using excitation wavelength of λ_Ex_=1300 and 1650nm respectively. Microglia in green, vessels labeled with Texas Red in grey, Glioma cells in magenta and the Third harmonic generation (THG) signal in cyan. Cellular resolution is reached throughout the entire z-stack as shown in the right panel. **E** Representative images of microglia cells in corpus callosum in the absence of glioma (ctrl) or in glioma infiltrated fiber tracks (GB+). Asterisks indicate representative 3D reconstructions of individual microglia as indicated in the lower panel for Sholl analysis. **F** Sholl intersection profile of microglia under GB free (black circles) conditions and under GB burden (white circles); note reduced arborization of microglia in GB affected volumes (averaged over n=4 mice with n(ctrl)=52 and n(GB+)=72 individual microglia cells) **G** Area under the curve (AUC) for the plot shown in E. (Averaged over n=4 mice; two sided *t*-test; *p*=0.03; *t*=2,828, *df*=6) **H** Schematic (upper panel) and representative images microglia cells and their processes for three different stages of GB progression within the CC: ctrl, sparse and dense (0%, <30% and >30% GB coverage). THG in grey first panel, Microglia in green and GB-cells in magenta. Subsequent time points (0 min, 5 min) were superimposed (Δ t = 5 min) to measure gained (green), lost (red) and stable (yellow) processes **I** Schematic of the concept of stable, loss and gained microglial fine processes for microglial motility analysis **J** Turnover rate of microglia as a measure of their surveillance activity (i.e. microglial motility), one-way ANOVA with Šídák’s multiple comparison test (J-N). (J) *F* (2, 34) = 21,28. Adjusted *p*-values: ctrl vs sparse p=0.0013; ctrl vs. dense *p*=0.0282; sparse vs dense *p*<0.0001; (H-K) n=13 ctrl, 14 sparse, 10 dense FOVs from individual experiments in n(ctrl)=5 mice, n(sparse)=7, n(dense)=3 and a total of n(ctrl)=89, n(sparse)=107 and n(dense)=122 microglia cells **K-M** Separation of microglia fine process turnover fractions into Stable(K), Loss(L) and Gain(M) fractions. (K) Stable fraction: *F*(2, 34)=15.59; adjusted p values: ctrl vs. sparse *p*=0.0103; ctrl vs. dense p= 0.0437, sparse vs. dense p<0.0001. (L) Loss fraction *F*(2, 34)=2; adjusted p-values: : ctrl vs. sparse p=0.9855; ctrl vs. dense=0.3181; sparse vs. dense p=0.1807 (M) Gain fraction F(2, 34)=12.22; adjusted p-values: : ctrl vs. sparse p=0.0217; ctrl vs. dense=0.0941; sparse vs. dense p<0.0001. **N** Number of individual microglia in a field of view for the three conditions. On average ctrl=6.85 ± 0.69 sparse 8,23 ± 0.83, dense=12.2± 1.14 microglia in field of view. n.s. not significant, * p < 0.05, ** p <0.01,*** p <0.001, **** p < 0.0001; Scalebars: 3 mm (A), 50 µm (B), 100 µm(C), 10 µm (D)

±1.3%, **Figure 4J**). In contrast, in densely GB-cell infiltrated CC microglial surveillance decreased by 25% (35.3 ±1.3%, **Figure 4J**). This downregulation of microglial surveillance during dense GB infiltration was mainly due to a marked increase in the stable fraction accompanied by a significant decrease of the gained processes fraction **(Figure 4K and 4L)**. On the contrary, in sparsely infiltrated CC, the initial increase of microglial TOR was fostered by a significantly increased gained fraction of microglial processes together with a marked reduction of stable processes **(Figure 4 K and 4L)**. Surprisingly, the loss fractions did not differ between the three groups **(Figure 4M)**. Moreover, we found a marked increase in microglial density in the densely GB-cell-infiltrated CC **(Figure 4N)**.

Our longitudinal analysis showed that GB-cells progressively populate the CC-adjacent cortex upon sustained GB-cell migration to the far infiltration zone in the contralateral hemisphere. In contrast to the CC, which is primarily composed of myelinated axon tracts (white matter), the cortex mainly consists of neuronal cell bodies and synapses (gray matter). We set out to investigate how microglial morphology and surveillance capacity were affected during progressive GB-cell infiltration of the contralateral cortex **(Figure 5A and 5B, upper left panel)**. In the presence of GB-cells, microglia displayed an amoeboid shape with reduced branching and morphological complexity **(Figure 5C)**. Sholl-analysis revealed a clear reduction in intersections of microglial branches in GB^+^ compared to GB^–^ cortical tissue **(Figure 5D and 5E).** We then decided to monitor microglia surveillance capacity upon encountering GB-cells. As above, we studied three distinct stages of GB-cell infiltration in the cortex; ctrl 0%, sparse <30% and dense >30% GB-cell infiltration levels **(Figure 5F upper panel; Videos S11-13)**. Microglial fine process turnover was measured **(Figure 5G)** for all three infiltrative stages. We did not find any differences in microglial surveillance between sparse infiltration and the absence of GB-cells **(Figure 5H)**. However, there was a drastic decrease of microglial motility of 41.6 % under higher levels of GB-cell infiltration (sparse 48.5 ±2.8% vs. dense 28.3% ±1.2% **Figure 5H**). This marked decrease in microglial motility was mostly due to a significant increase in immobile fine processes **(Figure 5I**) and a corresponding reduction in process expansion **(Figure 5J),** as well as retraction of microglial protrusions **(Figure 5K)**. In addition, the density of microglia was unchanged between ctrl conditions and sparse infiltration patterns **(Figure 5L)**. Only in densely infiltrated cortex, we found a marked 210% increase of microglial density (**Figure 5L and see example 5F**; sparse 8.1 ±1% vs. dense 25.2 ±3.7%).

**Figure 5.**
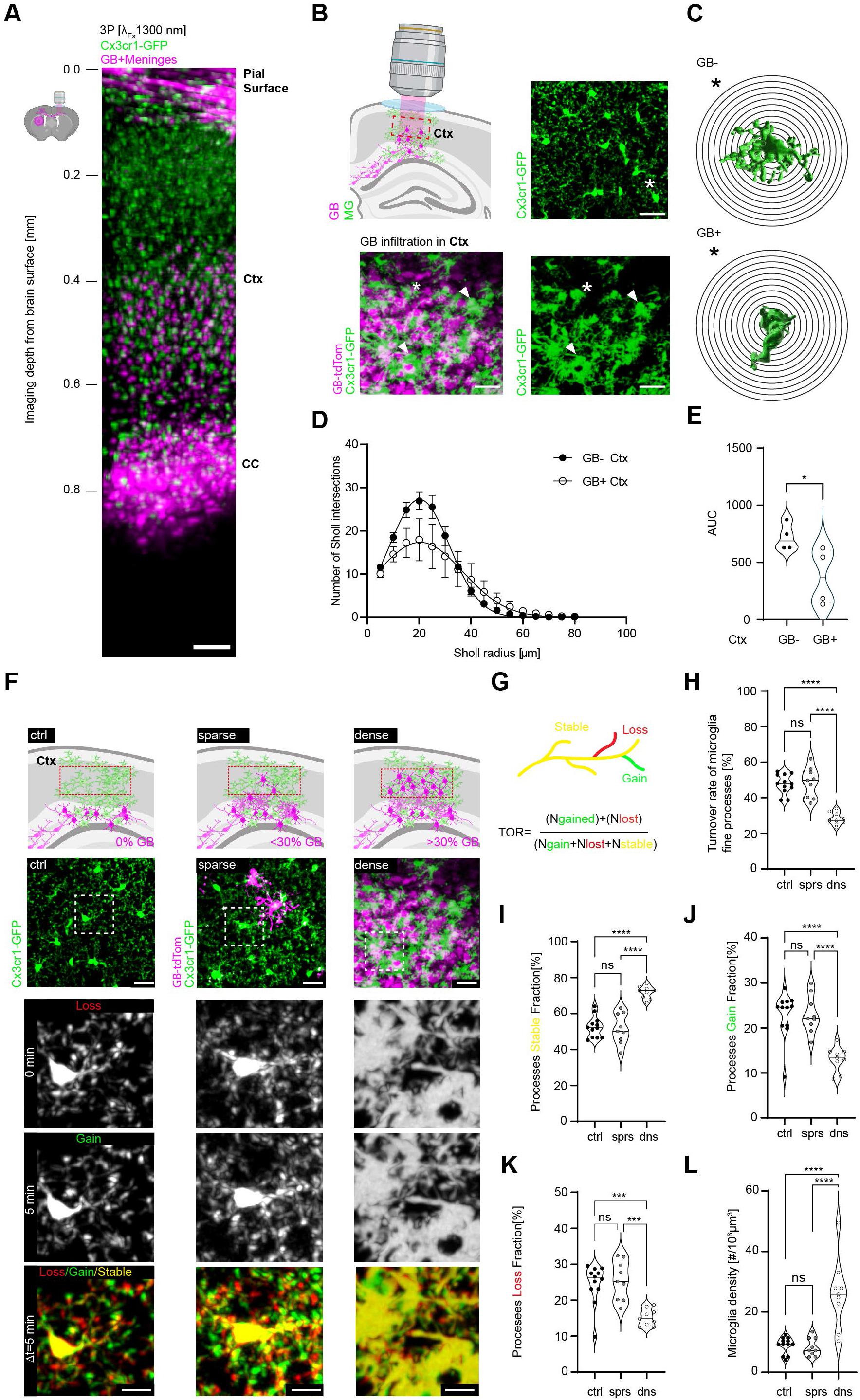
Fade out of microglia surveillance upon GB-cell infiltration of the cortex. **A** 3D reconstruction of a z-stack spanning approx. 900 µm through the entire cortex and the deep-seated corpus callosum on the contralateral side in respect to the bulky tumor/the injection side. MG (GFP) and GB-cells (tdTom) were visualized using λ_Ex_=1300 nm. In this example cortical GB-cell infiltration via the CC has populated the cortex up to ca. 400 µm below the pial surface. **B** Representative images of microglia cells in the cortex either in the absence of glioma (ctrl, upper panel) or in glioma infiltrated fiber tracks (GB+, lower panel). Asterisks indicate representative 3D reconstructions of individual microglia as shown on the right panel **C** for consecutive sholl-analysis (D, E). **D** Sholl profile of microglia under GB-cell free conditions (black circles) and under GB burden (white circles); Microglia under GB burden show reduced arborization (averaged over n=4 mice with n(ctrl)=159 and n(GB+)=151 individual microglia cells. **E** Area under the curve (AUC) for the plot shown in D. (Averaged over n=4 mice; two-sided *t*-test; *p*=0.045; *t*=2,523, *df*=6). **F** Schematic (upper panel) and representative images microglia cells and their processes for three different stages of GB-cell infiltration within the cortex: ctrl, sparse and dense (0%, <30% and >30% GB-cell coverage). Microglia in green and GB-cells in magenta. Subsequent time points (0 min, 5 min) were superimposed (Δ t = 5 min) to measure gained (green), lost (red) and stable (yellow) processes. **G** Schematic of the concept of stable, loss and gained microglial fine processes for microglial motility analysis. **H** Turnover rate of Microglia fine processes in percent for the three conditions ctrl, sparse and dense; one-way ANOVA with Šídák’s multiple comparison test (H-L). *F*(2,27)=33,68; Adjusted p-values: ctrl vs sparse p=0,99; ctrl vs. dense p<0.0001; sparse vs dense p<0.000. From (H-L) n=12 ctrl, n=9 sparse, n=9 dense FOVs from individual experiments in n(ctrl)=5, n(sparse)=4, n(dense)=3 mice and a total of n(ctrl)=104, n(sparse)=73 and n(dense)=227 microglia cells. **I-K** Separation of microglia fine process turnover fraction into Stable(K), Gain(L) und Loss(I) fractions. **I** Stable fraction: *F*(2, 27)=31,47; adjusted p values: ctrl vs. sparse *p*=0,98; ctrl vs. dense p<0.0001, sparse vs. dense p<0.0001. **J** Gain fraction *F*(2, 27)=17,19; adjusted p-values: ctrl vs. sparse *p*>0,99; ctrl vs. dense= p<0.0001; sparse vs. dense p<0.0001. **K** Loss fraction *F*(2, 27)=13,43; adjusted p-values: : ctrl vs. sparse *p*=0.96; ctrl vs. dense=0.0004; sparse vs. dense p=0.0003. **L** Number of individual microglia cells per identical imaged volume and FOV. *F*(2,27)=21,22; adjusted p-values: : ctrl vs. sparse *p*=0.996; ctrl vs. dense p<0.0001; sparse vs. dense p<0.0001; n.s. not significant, * p < 0.05,*** p <0.001,**** p < 0.0001; Scalebars: 100 µm (A), 20 µm (B, F upper panels), 10 µm (F, lower panels)

Taken together, 3P-imaging of the CC and deep cortical areas enabled us to discover a biphasic response in microglial surveillance activity upon progressive GB-cell infiltration. Furthermore, our data suggest differences in microglia behavior depending on their location in the white or gray matter.

### Identification of microglia subsets with distinct migration properties in relation to GB-cell infiltration

It has been proposed that microglia migrate to and accumulate at GB (near and far) infiltration zones.^58–60^ Microglia migration has been investigated in the near infiltration zone in a GL261 model with 2P *in vivo* imaging in superficial layers of the cortex.^23^ However, the kinetics and patterns of microglia migration at the far infiltration zone have not been studied. Using 3P microscopy, we imaged the same region of the somatosensory cortex and CC, contralateral to the electroporation side over several days at 24-hour intervals **(Figure 6A and 6B)**. We sampled large z-volumes spanning on average 1,000 µm through the entire cortical column and CC **(Figure 6C)**. This approach allowed us to investigate microglial migratory properties at different imaging depths and at different distances from invading GB-cells over several days. We found that microglia residing in GB-cell-free cortical areas remained stationary over time and did not exhibit significant migratory behavior **(Figure 6D, Video S14)**. In contrast, a marked increase in microglia migration was observed in close proximity to GB-cells migrating along the CC **(Figure 6E and 6F, Video S15)**. Notably, some microglia remained largely stationary even when situated near GB-cells over the imaging period (**Figure 6E**, white arrows, **Video S16**). Furthermore, microglia traveled 2.4-times larger distance when closer than 150 µm to the nearest GB-cell, in comparison to microglia further apart **(Figure 6F and 6G)**. Consistent with this finding, microglia in the non-infiltrated cortical areas exhibited lower variance in Euclidean distance to their six nearest neighbors compared to microglia in the CC at the far infiltration zone **(Figure 6H)**. It has been suggested that microglia, even when located in close proximity to each other, can respond differently to a given stimulus and exhibit divergent functional and transcriptional properties.^61,62^ To investigate the migratory responses of different microglia subsets, we conducted unbiased agglomerative clustering and principal component analysis on horizontal sub-volumes as depicted in Figure 6D and 6E **(Figure 6I)**. We identified four distinct subsets of microglia at the GB far infiltration zone. The primary biological components accounting for the variance in our data were: the distance of microglia to the nearest GB-cell **(Figure 6J)**, the variance in Euclidean distances between neighboring microglia **(Figure 6K**), and their traveled distances **(Figure 6L)**. Microglia in Cluster 1 (C1) were in close proximity to the tumor within the CC and exhibited a relatively uniform distribution, but showed increased travelled distances (**Figure 6M**). C2 microglia were also located near the neoplastic compartment, but were distributed heterogeneously and display minimal migration (**Figure 6M**). C3 comprises microglia that were located further away from the tumor and were likely cortical microglia, characterized by a relatively uniform spatial distribution and short travelled distances (**Figure 6M**). Finally, C4 contained microglia situated at the GB far infiltration zone, with a homogeneous distribution pattern and moderate-to-small travelled distances (**Figure 6M**). To further investigate microglia subsets at the far infiltration zone, we analyzed migration patterns along the z-axis of our microscopy volume and applied agglomerative clustering to categorize these subsets accordingly **(Figure 6N)**. We identified four distinct clusters of microglia, which accounted for most of the variance observed in our data **(Figure 6N)**. Consistent with our previous findings, C1 had the shortest distance to the nearest GB-cells **(Figure 6O)**, whereas C4 represented cortical microglia **(Figure 6O)**. C1 also exhibited a significantly higher average traveled distance compared to the other clusters **(Figure 6P)**. Notably, only C2 demonstrated directed movement toward GB-cells as indicated by their changes in distance to the nearest GB-cells **(Figure 6Q)**. This cluster represented microglia at the GB border that are relatively close to the tumor, but not yet in direct contact **(Figure 6P)**. Finally, microglia located within tumor infiltrated tissue of the contra-lateral hemisphere exhibited rather random, non-directed movement (**Figure 6Q**).

**Figure 6.**
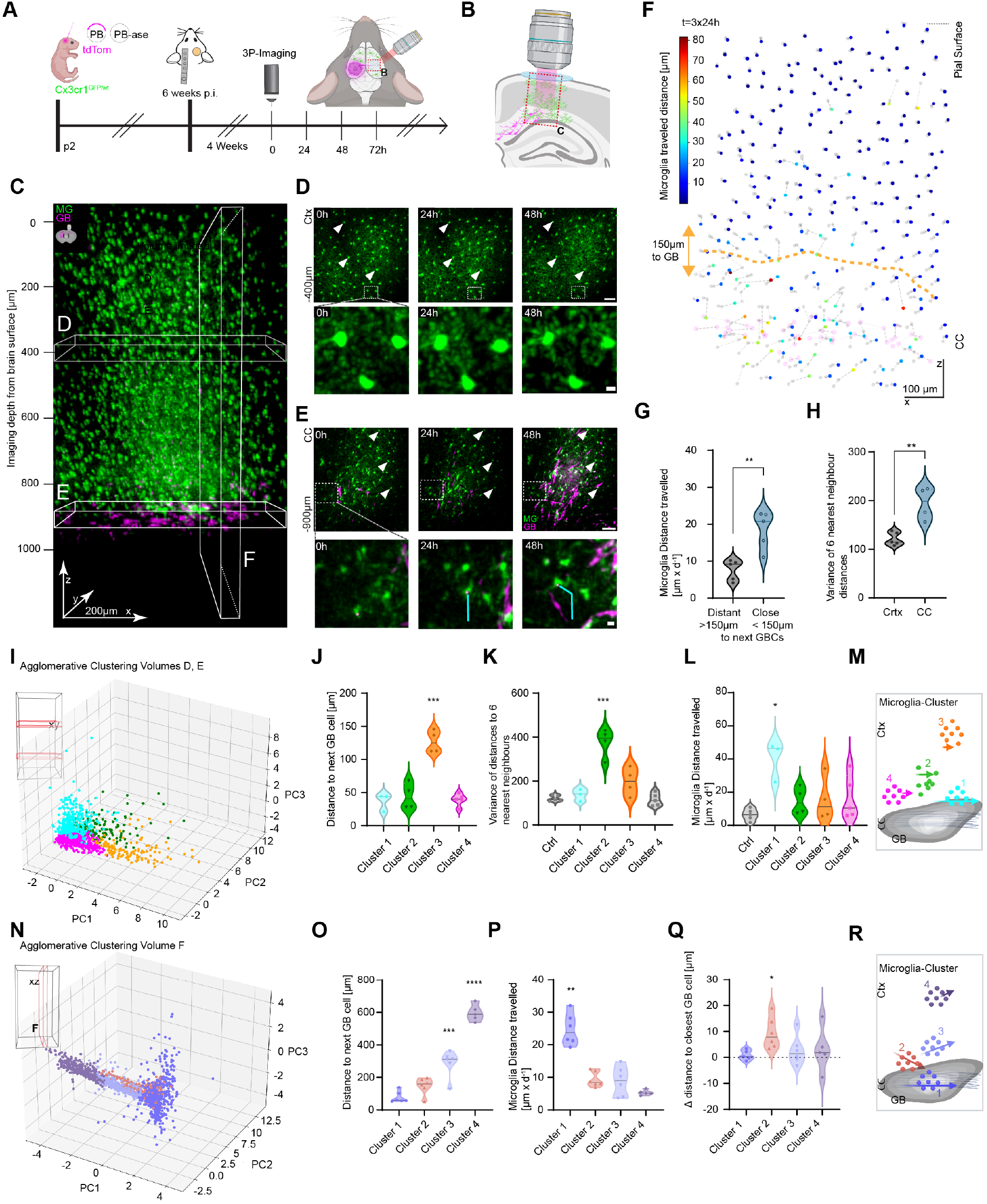
Dimensional reduction reveals subsets of microglia differing by their migration properties towards invading tumor cells. **A** As a recipient we use the Cx_3_Cr-1 ^GFP/wt^ mouse line. At P2 the CRISPR/Cas plasmid mix (sgRNAs against Nf1, Pten and p53 and a Transposom system for fluorescent visualization) is injected into the lateral ventricle with consecutive electroporation and cranial window surgery as shown previously (Fig.1,2). About 3 months after tumor induction intravital 3P microscopy is performed with a 24h-interval between imaging sessions. **B** 3P imaging allows non-invasive detection of early stages of GB-cell infiltration via the CC **C** 3D side projection of MG (green) through the entire cortex and into CC harboring migrating GB-cells (magenta) **D** Cortical microglia in the absence of GB-cells imaged over 3 days (24h-interval) with a distance of approx. 500 µm to the GB-cell infiltrated CC. Arrows exemplary mark non-migrating, stationary MG cells. Zoom in lower panel shows how little dislocation microglial somata display under unperturbed conditions. **E** Microglia from the same imaging session as in D but in CC. The example depicts early infiltration, with single individual cells migration via CC at day0 to rather dense infiltration 48hours later. Asterisks mark stationary microglia. The lower panel shows a microglial cell marked with a sphere, which is migration in response to the onset of GB-cell infiltration in the CC (see cyan trajectory). **F** Visualization of a xz-dimension sub volume as indicated in C in which individual microglia are displayed as dots with a color coding from blue to red depending on their traveled distances over 3 days with a 24h-interval. The migration direction of each cell is displayed as a dashed line. GB-cells are colored in light magenta within the CC approx. 900 µm below the pial surface. The orange dashed line shows a 150 µm perimeter around the infiltrating GB-cells in which microglia are likely to react to GB-cell presence. **G** Microglia traveled distances par day at >150 or <150 µm range to next GB-cell, unpaired *t*-test, two tailed, *p*=0,0031*, t*=4,186 *df*=8 from n=5 mice **H** Comparison between the variances of MG and their respective 6 nearest neighbors in Ctx and CC. Cortical unaltered microglia show a much more homogenous distribution in the tissue indicated by a low variance of distances to 6 nearest neighbors. Unpaired *t*-test, two-tailed*, p*=0.0028*, t*=4.49*, df*=7 from n(Ctx)=5 and n(CC)=4 mice. **I** 3D principal component analysis (PCA) displaying 4 distinct states of microglia obtained from different xy-subvolumes. **J** Distances to next GB-cells between Clusters 1-4, one-way ANOVA with Tukey’s multiple comparison test (J-L), *F*(3,12)=33.33 **K** Microglia variances between 6 nearest neighbors between clusters, ctrl is complete GB-cell free cortical tissue as internal reference, *F*(4,16)=26.23 **L** Comparison of MG traveled distances between the 4 Clusters and the internal control, *F*(4,16)=5,06 **M** Schematic visualization of the 4 clusters with respect to their relative distance to the next GB-cells, their variance in distances amongst each other and their respective travelled distances, depicted by the arrow length. **N** 3D PCA showing 4 distinct states of MG found in the data obtained from xz-subvolumes. **O** The major determinant for cluster separation is the distance of microglia to the next GB-cells, one-way ANOVA with Tukey’s multiple comparison test (O-Q), *F*(3, 19) = 83,11 **P** These clusters also separated nicely by their respective travel distance of microglia, *F*(2,015, 9,405) = 46,85 **Q** The delta distance to closest GB-cell reflects the relative movement to each other between microglia and GB-cells. Although the multiple comparison test did not show significant differences between clusters, *F*(3, 13) = 3,55, there is at least a strong tendency between CL1 and CL2 (adjusted p-value = 0.068). **R** Schematic visualization of the 4 clusters from xz sub volumes with respect to their distance to the next GB-cells, respective travelled distances and their relative movement to the GB-cells depicted by the arrow length and orientation. n.s. not significant, * p < 0.05,** p <0.01, *** p <0.001,**** p < 0.0001; Scalebars: 50 µm (D,E, OVs), 10 µm (E, lower panel), 5 µm (D, lower panel)

In summary, we developed a 3P-imaging approach based on a clinically relevant autochthonous model that provided insights into the respective GB-cell and microglia *in vivo* behaviors at the GB far infiltration zone, with implications for our understanding of tissue surveillance and tumor invasion.

## Discussion

In this study we reveal the spatio-temporal dynamics of microglia interaction with GB-cells in an autochthonous immune-competent GB mouse model at the far infiltration zone. Performing 3P-imaging, we found a biphasic response of microglial surveillance upon increasing GB-cell infiltration. Moreover, microglia at the far infiltration zone form a heterogeneous population with diverging migratory properties in proximity to invading GB-cells.

The diffuse nature of GBs along with the aberrant immune-TME represents key barriers to existing treatments of this devastating disease. Despite decades of intense research, our understanding of how innate immune cells detect and react to GB invasion remains incomplete. It is therefore not surprising that immunotherapies have failed to replicate preclinical data in clinical trials.^63,64^ One reason for this discrepancy may be the inadequate immune responses associated with rapid tumor growth in common GB-models, which often are based on xenotransplantation of established tumor cell lines. Furthermore, until recently our ability to capture *in vivo* behaviors of tumor and immune cells at the GB infiltration areas in a living and intact brain tissue has been hampered by insufficient imaging methods preventing deep brain imaging at one millimeter depth. To overcome these shortcomings, we used an autochthonous mouse model of Mes GB,^3,36^ which recapitulates key aspects of human GB pathology and performed 3P intravital microscopy in deeply located brain structures, like the CC.

First, we investigated the relationship between TMs, a hallmark feature of IDH^WT^ GB, ^65,66^ and GB-cell migration, which has been studied mainly applying two-photon imaging in superficial cortical areas.^9,67^ TMs are up to 500 µm long, thin protrusions of the plasma membrane, integrating GBCs into a cooperative multicellular functional network.^52,68^ These networks convey to resilience against multimodal treatment regimens.^65,66,69^ Within these networks GBCs display a highly ramified morphology with many TMs, whereas infiltration of non-affected brain parenchyma is driven by unconnected GBCs with less TMs, as shown in different PDX models.^9^ In a recent study, 3P-microscopy has been applied to describe TM dynamics of GB-cells inside the CC, in an immunodeficient PDX model of GB in proximity to the tumor bulk.^70^ Beyond that, our data derived from an immune-competent GB-model, in which we interestingly found a large fraction of fast-migrating GB-cells in the CC, in contrast to GB-cells in the cortex that were primarily stationary. In addition to previous findings, our data underscore that TMs inversely correlate with GB-cell migration at the far infiltration zone in the CC, as in closer proximity to the tumor bulk. It is tempting to speculate that TMs might, besides their role in network function, also act as basic cellular components anchoring GB-cell in the parenchyma thus limiting GB-cell mobility. According to this, higher numbers of TMs might also be a predictive factor for their network integration ability. Therefore, while targeting TMs represents a potentially effective therapeutic target, as we and others have reported,^71,72^ release of tumor cells from such networks may enhance their migration capacity, a phenomenon that should be further investigated.

We also discovered that GB-cells travel faster within the CC than in the cortex. It is possible that significant differences for instance in the composition of the extracellular matrix between the two anatomical compartments contribute to this phenomenon. Myelin is enriched in the CC and contains proteins that inhibit cell migration such as Nogo-A, myelin-associated glycoprotein, and oligodendrocyte myelin glycoprotein.^73,74^ Interestingly, GB-cells are rich in proteases including various metalloproteinases, thus able to cleave these inhibitory molecules from the myelin sheaths of the CC. These GB-cell properties potentially make the CC a migratory scaffold by conditioned remodeling of the extracellular matrix.^75^ Whether and how these molecular mechanisms play a role for different migration speeds, remains to be determined in the future.

Within the non-neoplastic TME compartments, microglia represent approximately 30% of tumor bulk and have been shown to closely interact with GB-cells, although their pro- or anti-tumor roles in this disease remain not fully understood,^12,15^ potentially because of their reported heterogeneity.^76,77^ Another reason for our incomplete understanding is that it is often based on extrapolation of *ex vivo*/*in vitro* experiments or relies on preclinical models based on transplanted tumor cell lines in immunocompromised recipients. One of the first studies aiming to bridge the knowledge gap about GB-microglia interaction dynamics *in vivo*, based on 2P microscopy in living mice transplanted with the GL261 tumor cell line in the cortex.^25^ Microglia acquired an amoeboid shape within 36h after tumor seeding. Another report using the same model revealed similar morphological changes of microglia and described migration properties of microglia in proximity to the tumor bulk.^23^ Two further studies measured microglial process dynamics in a bulky orthotropic GB implantation model and a xenotransplantation model, respectively.^24^ However, longitudinal measurements of microglia migration and fine process motility in a GB model have not been performed in CC and adjacent cortex at the far infiltration zone up to now. We recently established 3P *in vivo* imaging to measure microglial fine process motility more than a millimeter deep in the prefrontal cortex under normal physiologic conditions.^42^ Here, we applied and adapted this 3P approach to investigate the dynamics of microglia in relation to GB-cells at the far infiltration zone. We identified a marked loss of cellular complexity of microglia. This is consistent with various studies showing de-ramification of microglia under GB burden in different GB mouse models and human GB samples.^25,78–82^ Furthermore, microglia cell numbers within GB tissue were increased in human GB sections consistent with our findings and multiple studies in mouse models showing elevated numbers of microglia in GB infiltrated tissue.^12^ Strikingly, we observed that microglial process motility initially increased upon sparse infiltration of GB-cells into the CC. Later, in densely infiltrated GB, microglial surveillance faded. This biphasic microglial response suggests an initial antitumor reaction, with microglia attempting to limit tumor spread. However, when GBs progress to dense infiltration, the subsequent decrease in microglial activity suggests acquisition of aberrant disease-associate immunosuppressed states, as observed in other brain pathological conditions.^17,83,84^ These GB-associated microglia phenotypes have been for instance linked to GB-cell secreted IL-10, TGF-b CXCL12, GDNF, EVs transporting miRNA21 and potentially also lipid droplets.^12,85,86^ A long-lasting enhancement of these immune escape mechanisms of GB-cells is predicted to be crucial for disease progression and successful infiltration. Our measurements of microglial motility in GB-infiltrated cortex however, did not replicate the aforementioned biphasic microglia response in the CC, but they pointed at a more general decrease of microglial motility in densely infiltrated GB tissue. These data suggest fundamental differences in microglia behaviors between CC and cortex potentially related to different micro-environments in white and gray matter.^87^ ^88,89^

Our immune-competent somatic model allowed us to investigate the dynamics of microglia/tumor cell interactions.^70^ In this respect, it has been previously proposed that microglia migrate to the far GB infiltration zone,^23,60^ but the underlying kinetics have not been investigated *in vivo*. Consistent with previous studies reporting only minimal migration of microglia in cortical parenchyma in the absence of pathological stimuli,^7,56,90,91^ we also barley observed microglia migration within the non-infiltrated, undisturbed cortex. We found that microglia migrated towards GB-cells within a distance of approximately 150 µm at the far GB infiltration zone. This fits well to previous studies in different lesion models observing a recruitment radius of microglia between 100 – 200 µm.^92^ Which mechanisms contribute to microglia recruitment to GB-cells at the far infiltration zone, remains to be discovered in the future.

Finally, single-cell and spatial transcriptome profiling has pointed at the molecular heterogeneity of microglia within GB^3,60,76,93^ and in other disease models.^94^ In this respect, our findings suggest functional heterogeneity with respect to how tumor cells (invasion) and microglia (surveillance and migration) interrelate in the context of an immune-competent preclinical model. This work has also broader implications for our understanding of how the brain innate immune system reacts to emergence of pathology. This in turn could lead to discovery of new actionable mechanisms to enhance microglia ability to preserve tissue homeostasis during aging, in cancer and other brain pathological states.

### Limitations of the study

Our work indicates a biphasic early response mechanism of microglia upon GB-cell contact. Although it is tempting to speculate that these changes in microglial motility represent an initial antitumor response, their molecular features associated with these functional changes remain unknown. Furthermore, the replacement of one fractalkine receptor allele with GFP (i.e. Cx3cr1^GFP/WT)^ may influence our findings. However, it has been shown previously that Cx3cr1 heterozygosity does not have any effect on the migration capacities of tumor associated microglia.^95^ Finally, it is not clear if our findings — both the diffuse GB infiltration and the microglial responses — can be generalized to other GB subtypes, as we focused on the mesenchymal subtype in this project. This remains an aspect for future investigation.

## RESOURCE AVAILABILILTY

### Lead contact

Further information and requests for resources and reagents should be directed to and will be fulfilled by the lead contact, Felix Nebeling.

### Materials availability

All unique reagents generated in this study are available from the lead contact with a completed material transfer agreement.

### Data and code availability

The authors declare that the data supporting the findings of this study are available within the paper, the methods section, and Supplementary data files. Data are available open access from DRYAD.

## ACKNOWLEDGEMENTS

FN received funding from the Mildred-Scheel School of Oncology Cologne-Bonn. This work was supported by the DZNE, and grants to MF and PS by the European Union ERC-CoG (MicroSynCom 865618, H3.3Cancer 616744). MF received funding from the German research foundation DFG (SFB1089 C01, B06; SPP2395) and is a member of the DFG excellence cluster ImmunoSensation2. This work was also supported by the iBehave and CANTAR network to MF, FN, PS, MH (funded by the Ministry of Culture and Science of the State of North Rhine-Westphalia; the funders had no role in study design, data collection, and interpretation, or the decision to submit the work for publication). PS received funding from the Wilhelm Sander Stiftung, the Worldwide Cancer Research Fund (WCRF) and the German Cancer Aid-funded THUNDER program. PS and DB thank the Helmholtz-Gemeinschaft Aging and Metabolic Programming (AMPro) Consortium (to PS, DB). We would like to thank present and former members of PS group involved in this work, along those in the group of DB, the Core Research Facilities, especially the Light microscopy facility (LMF), Services at DZNE, and the DZNE Animal Research Facilities for continuous support through the project. PS is recipient of an Honorary Professorship at University College London (2023-2027). We thank Ute Heuser-Figgemeier and Alexandra Brüggemann from the Institute of Neuropathology, University Hospital Bonn, Bonn Germany for their support with tissue processing. Thanks to Pinzhang Lu for technical assistance. We thank Julia Steffen, Andrea Baral and Yvonne Schleehuber from the Neuroimmunology and Imaging group (DZNE) for providing technical assistance. Additionally, we would like to thank P. Thevenaz for the development of the ImageJ plugins “stackreg”. Figures were prepared with Illustrator CS5 Version 15.0.1 (Adobe) and created with with BioRender.com (Details see below)

## AUTHOR CONTRIBUTIONS

FN conceptualized the experiments, collected data, analyzed data, designed the figures and wrote the manuscript. FF provided technical expertise and support for 3P-imaging. FM helped analyzing microlgial motility and carried out image registration. MM helped analyzing microglia migration and performed PCA. NG analyzed microglia migration. LF selected and imaged the human GB samples. SL helped with histological analysis. AD technically assisted initial electroporation. MS wrote animal experimental protocol and obtained the license. SF provided technical support for the 3P-microscope. DB provided funding and reagents. TP provided neuropathological samples and expertise. SP provided the plasmids and helped with technical expertise. MH provided resources and consulting on conceptualization. UH provided conceptualization and translational input and helped writing the manuscript. PS provided conceptualization, technical expertise and helped writing the manuscript. MF provided technical expertise, analyzed data, conceptualized experiments, wrote the manuscript. MF, PS and UH supervised the project.

## STAR * METHODS

### KEY RESOURCES TABLE

| **Reagent or Resource** | **Source** | **Identifier** |
| --- | --- | --- |
| Chemicals, Peptides, and Recombinant Proteins |  |  |
| Buprenorphine | Reckitt Benckiser |  |
| Isoflurane | Virbac | Vetflurane® |
| Metamizol-Natrium 1 H<sub>2</sub>O | Serumwerk Bernburg | Metapyrin<br>CAS: 68-89-36 |
| Ketamin | Medistar | Ketamin 10% Inj.-Lsg. |
| Xylacin | Bayer | Rompun® |
| Texas Red™ | ThermoFisher | Cat# 1864 |
| Ki-67 Antibody | Abcam | Cat#16667 |
| CD11b Antibody | BD Biosciences | Cat#565080 |
| CD44 Antibody | BD Biosciences | Cat#558739 |
| CD68 Antibody | Biolegend | Cat#916104 |
| PBase<br>bicistronic construct with<br>SpCas9, PiggyBase under<br>hGFAPmin promoter | Simona Parrinello | N/A |
| PBvector<br>tdTomato fluorescent<br>reporter, sgRNAs for Nf1,<br>Pten and Trp53. | Simona Parrinello | N/A |
| Fastgreen | Sigma | Cat#P6148 |
| Critical commercial assays |  |  |
| Plasmid Mega Kit | Qiagen | Cat#12181 |
| Experimental Models: Organisms/Strains |  |  |
| Mus musculus:<br>C57BL/6J | The Jackson Laboratory | Strain #: 0664 |
| Mus musculus:<br>B6.129P-CX3CR1<sup>tm1Litt</sup>/J | The Jackson Laboratory | Strain #: 5582 |
| Software and Algorithms |  |  |
| Fiji | https://fiji.sc/ | N/A |
| Imaris | https://imaris.oxinst.com/ | N/A |
| Python | https://www.python.org/ | N/A |
| GraphPad Prism | https://www.graphpad.com/ | N/A |
| Zen microscope software | https://www.zeiss.com/microscopy/de/home.html | N/A |
| Biorender | https://biorender.com/ | N/A |

## METHOD DETAILS

### Animals

Mice were group-housed and separated by gender with a day/night cycle of 12 hr. Water and food was accessible *ad libitum*. All experiments were performed according to animal care guidelines and approved by the Landesamt für Natur, Umwelt und Verbraucherschutz of North Rhine-Westphalia (Germany) (#81-02.04.2018.A148). C57BL/6J (Stock:000664) were obtained from Jackson Laboratory. CX3CR1-GFP knock-in mice carry the international name: B6.129P-CX3CR1^tm1Litt^/J (Stock: 5582) Jackson Laboratory. The mouse line expresses eGFP in monocytes, dendritic cells, NK cells, and microglia in the brain. The *Gfp* gene is introduced as a knock-in into the allele of the *Cx3cr1* gene (chemokine C-X3-C motif receptor 1). These mice were first described by Jung et al.^55^

### *In vivo* electroporation

Tumor induction was performed as described before.^3^ In short, two separate plasmids were injected into the left ventricle of mice at postnatal day2 after 60-90 sec of cold-shock immobilization. One plasmid contained the gRNAs for Nf1, Pten and P53 alongside the tdTomato fluorescent reporter under the hminGFAP reporter.^3^ The Other carried the episomal SpCas9 and the transposase for stable tdTom expression in the transfected NSCs and their progeny. The PiggyBase (0.5 μg/μl) and PiggyBac vectors at a molar ratio of 1:1 were diluted in TE buffer together with 0.1% fast green dye (Sigma) for visual tracing of the injection. Electroporation was performed with 5 square pulses with 50msec/pulse at U=100V and with 850 msec intervals.

### Cranial window surgery

Cortical window surgery carried out four weeks prior to imaging. Mice were anesthetized with an i.p. injection of ketamine/xylazine (0.13/0.01 mg/g body weight). As an analgetic mice received buprenorphine (0.1 mg/kg; subcutaneously (s.c.), Reckitt Benckiser) shortly before surgery. The body temperature was maintained with a heating pad at 37°C. After fixation in a stereotaxic frame, the skin was removed under aseptic conditions. Using epifluorescence, the location of the bulky tumor was visualized below the left somatosensory cortex and accordingly and a craniotomy above the right somatosensory cortex (4 mm diameter) was performed with a dental drill. The dura was carefully removed. The brain surface was rinsed with sterile saline, and #1 coverslips (4 mm diameter) were sealed into the craniotomy with dental cement. Post surgery, mice received buprenorphine (0.1 mg/kg s.c.) three times daily and Metamizol (200 mg/kg) applied to the drinking water for 3 consecutive days. For head-fixation during *in vivo* imaging a headpost (Luigs & Neumann) was cemented adjacent to imaging window. For analgesia, buprenorphine (0.1 mg/kg s.c.) was injected three times daily and Metamizol was (200 mg/kg) was applied to the drinking water for 3 consecutive days.

### Microscope

The microscope has recently been described in detail by our group.^42^ In short, 3P-excitation is achieved with a wavelength-tunable excitation source from Spectra-Physics (NOPA, Spectra-Physics) pumped by a femtosecond laser (Spirit, Spectra-Physics). The laser repetition rate is maintained at 2 kHz. The 2P excitation source is a Ti:sapphire laser (Chameleon Ultra II, Coherent). The laser repetition rate is at 80 MHz. Images are taken with a multiphoton microscope built of a Thorlabs microscope base (EMB100/M) and primary scan path optics with 900-1900 nm coatings. Two GaAsP photomultiplier tubes (PMT2100, BDM3214-3P, Thorlabs) are used for non-descanned detection. ThorImage (Version 4.3, Thorlabs) is used to control imaging parameters. A 25x water immersion microscope objective with the numerical aperture of 1.05 (XLPLN25XWMP2, Olympus) is used. Green and red signals are separated by a 488 nm dichroic mirror (Di02-R488, Semrock) and 562 nm dichroic mirror (FF562-Di03). Then the eGFP and third harmonic generated (THG) signals are further filtered by a 525/50 nm band-pass filter (FF03-525/ 50, Semrock) and 447/60 nm (FF02-447/ 60, Semrock) band-pass filter, respectively. Lateral movements are done with the 2D stepper motor (PLS-XY, Thorlabs), z-stacks are acquired using the built-in z-axis motor or a piezo focus device (PFM450, Thorlabs).

### In vivo imaging

Mice were anesthetized with isoflurane (1% in 0,7l O2/min.) and head-fixed under the microscope on a heating pad at 37°C. Deep overview *in vivo* z-stacks were recorded with depth increments of 7 µm and 0.65 µm/pixel resolution and 1.49 µs pixel dwell time. Z-stacks were spanning 900-1100 µm from the pial surface through the somatosensory cortex and CC. For visualization of CC and tdTom+ GB-cells excitation wavelengths of λEx=1650nm was used. Microglia (Cx3cr1-GFP) and GB-cells (tdTomato) were co-imaged using λEx=1300nm. For the measurement of microglial fine process motility, z-stacks of 166.4x166.4x130-150 µm were acquired at 5 µm depth increments, 0,162 µm/pixel, 0.99 µs or 1.49 µs dwell time and with a 5-minute time-interval between z-stacks for a period of 25-30 minutes.

### Histology

Mice were deeply anesthetized with an i.p. injection of ketamine/xylazine and transcardially perfused with saline followed by either 4% paraformaldehyde (PFA) or directly fresh frozen at -80°C after extracting the brain. For visualization of the tumor anatomy and infiltration, brains were cut into 70 µm thick coronal slices. The tdTomato from the Piggybac system remained clearly visible without counterstaining. During tissue permeabilization (0.5% Triton-X100, 1h), DAPI was added to the free floating sections for 30 minutes (1:10000). For the visualization of CD44, Cd11b and Ki67, coronal 70 µm slices were permeabilized (0.5% Triton-X100, 1h) and subsequently incubated with the respective antbodies (CD44, CD11b, Ki67) in a blocking reagent (4% normal goat serum, 0.4% Triton 1%, and 4% BSA in PBS) over night at room temperature. After washing the samples three times with PBS, secondary antibodies were administered (Alexa Fluor 647, A21235, Alexa Fluor 594, A11058, Alexa Fluor 488, A48262) in 5% normal goat serum/BSA and incubated for 2h at room temperature. During the last 15 minutes of incubation DAPI was added (5mg/ml, 1:10000). Afterward slices were washed three times with PBS, mounted with Dako Mounting Medium, and covered with a glass cover-slip.

### Confocal microscopy

For the visualization of GB-cell infiltration patterns LSM800 microscope (Zeiss) was used. Entire coronal PFA-fixed brain slices of 70 µm thickness were imaged using a 20x Objective, with a lateral resolution of 0.624 µm/pixel. DAPI-(EX: G 365, Dichroic: FT 395, EM: BP 445/50), tdTomato (EX 561/10, Dichroic: 573, EM: 600/50) filter-sets were used.

### Human Samples

Human pretreatment GB samples were derived from Patients within the GLORIA Trial^96^ (SNOXA12C401, 2018-004064-62, NCT04121455) including patients with newly diagnosed, histologically confirmed, supratentorial CNS WHO grade 4 GBs, MGMT promoter methylated.

## QUANTIFICATION AND STATISTICAL ANALYSIS

### Microglial motility analysis

In general, microglial motility was analyzed as described before.^56,57^ In short, stacks were initially rigidly registered using subpixel image registration by cross-correlation^97^ provided by the open-source Python image processing library Scikit-image.^98^ Individual stacks were then median filtered and individual microglia were identified by scrolling through the stacks at each time point and cropped using ImageJ. Z-stacks spanning 35 μm in depth were isolated from the original z-stack. Stacks were registered by applying the ImageJ ‘StackReg’ plugin (ref). For all time points, the average intensity projections was calculated, resulting in a 2D visualization of microglia and their fine processes. Individual 2D Images were merged into a stack. Time points were pseudo colored in red and green. Red areas account for lost, green for gained and yellow for stable microglial fine processes. The turnover rate (TOR) of individual microglia processes was calculated as the number (absolute pixel value) of lost, Nlost (red), and newly gained, Ngained (green) pixels divided by the sum of all pixels within a determined region of interest (ROI). Building up on these manual analyses we implemented a custom analysis pipeline for automated quantification written in Python as described before ^42^; in summary, we calculated the temporal variation *ΔB(ti)* of binarized images by subtracting the binarized image *B(ti+1)* at time point *ti+1* from the binarized image *B(ti)* at time point *ti*: *ΔB(ti) = 2×B(ti+1)-B(ti)* for *i=0, 1, 2, …, N-1*, where *N* is the total number of all time lapse time points. Pixels in *ΔB(ti)*, that have the value 1, were categorized as stable pixels, whereas pixels with the value -1 were categorized as gained pixels, and pixels with the value 2 as lost pixels. The microglial fine process motility was then assessed by calculating the turnover rate (TOR) as the ratio of the number of all gained pixels *Ng(ti)* and all lost pixels *Nl(ti)* divided by the sum of all pixels: *TOR(ti) = (Ng(ti)+ Nl(ti)) / (Ns(ti) + Ng(ti)+ Nl(ti)),* where *Ns(ti)* is the number of all stable pixels. The average turnover rate 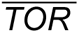 was calculated by averaging TO*R(ti)* over all *N-1* time lapse time points.

### Tumor microtubes

Cellular protrusions were classified as previously described ^9^. TMs were defined as between 1.25-3 µm thick and with a length of at least 10 µm.

### Principal component analysis

For the clustering analysis, tools from pythons scikit-learn library were used. First, the data was standardized using scikit-learns StandardScaler algorithm. Principal component analysis was performed in order to decrease the number of feature dimensions. The first four principal components (PC) accounted for over 75% of the variance. Agglomerative clustering was performed, using the Euclidean metric and the linkage criterion ‘ward’.

### Statistics

Quantifications, statistical analysis, and graph preparation were carried out using GraphPad Prism 9 (GraphPad Software Inc, La Jolla, CA, USA). To test for normal distribution of data, D’Agostino and Pearson omnibus normality test was used for sample sizes of n>6 and the Shapiro-Wilk normality test for n<6. Statistical significance for groups of two normally distributed data sets paired or unpaired two-tailed Student’s t-tests were applied. If no normal distribution was evident, Mann-Whitney test for groups of two was used. One-way ANOVA with Tukey’s or Bonferroni’s multiple comparison test were performed on data sets larger than two, if normally distributed. For comparison of more than two not normally distributed data sets the Kruskal-Wallis test was performed with Dunn’s correction for multiple comparisons. If not indicated differently, data are represented as mean ± SEM.

## SUPPLEMENTAL INFORMATION

**Video S1**

Animated coronal view of an entire brain showcasing the diffuse GB-cell infiltration patterns in the autochthonous model

**Video S2**

Z-Stack spanning the entire cortical column and corpus callosum. THG(green) and tdTom+ GB-cells(magenta) excited at λEx=1650nm

**Video S3**

GB-cells migrating via the CC visualized using 3P-excitation at λEx=1650nm and recording the THG signal (green) and GB-cells (tdTomato, magenta) over 210 minutes.

**Video S4**

3D volume showing THG(green) and GB-cell (magenta) density increase over 3 consecutive days, 24-h interval

**Video S5**

Z-Stack spanning the entire cortical column and corpus callosum. Microglia (green) and texasRed Dextran labeled vessels(grey) excitation at λEx=1300nm, THG (cyan) Signal λEx=1650nm, tdTom+ GB-cells (magenta) are excitable at both wavelengths,

**Video S6**

3D volume rotating, showing Microglia (green), texasRed Dextran labeled vessels (grey), tdTom+ GB-cells (magenta) and the THG Signal (cyan)

**Video S7**

Microglial (green) Motility within the CC in the absence of infiltrating GB-cells, example recorded over 25 minutes

**Video S8**

Microglial (green) Motility within the CC during sparse infiltration of GB-cells (magenta) at the far infiltration zone recorded over 30 minutes

**Video S9**

Microglial Motility within the CC during sparse infiltration of GB-cells (magenta) at the far infiltration zone example 2, recorded over 30 minutes

**Video S10**

Microglial Motility within the CC during dense infiltration of GB-cells (magenta) at the far infiltration zone recorded over 30 minutes

**Video S11**

Microglial motility (green)in the cortex in the absence of infiltrating GB-cells, recorded over 25 minutes

**Video S12**

Microglial (green) Motility within the cortex during sparse infiltration of GB-cells (magenta), recorded over 35 minutes

**Video S13**

Microglial Motility within the cortex during dense infiltration of GB-cells, recorded over 25 minutes

**Video S14**

Insignificant microglia (Green) migration in GB-free cortical tissue, 4 days recording, 24h-intervals

**Video S15**

Elevated migration of microglia (gree) during GB (magenta) infiltration at the far infiltration zone in CC, 3 days recording, 24h-intervals

**Video S16**

Microglia migration (green) during GB (magenta) infiltration at CC/Cortex border, 4 days recording, 24h intervals

## REFERENCES

1. Ostrom, Q.T., Price, M., Neff, C., Cioffi, G., Waite, K.A., Kruchko, C., and Barnholtz-Sloan, J.S. (2023). CBTRUS Statistical Report: Primary Brain and Other Central Nervous System Tumors Diagnosed in the United States in 2016-2020. Neuro Oncol 25, iv1-iv99. 10.1093/neuonc/noad149.

2. Chinot, O.L., Wick, W., Mason, W., Henriksson, R., Saran, F., Nishikawa, R., Carpentier, A.F., Hoang-Xuan, K., Kavan, P., Cernea, D., et al. (2014). Bevacizumab plus radiotherapy-temozolomide for newly diagnosed glioblastoma. N Engl J Med 370, 709–722. 10.1056/NEJMoa1308345.

3. Garcia-Diaz, C., Poysti, A., Mereu, E., Clements, M.P., Brooks, L.J., Galvez-Cancino, F., Castillo, S.P., Tang, W., Beattie, G., Courtot, L., et al. (2023). Glioblastoma cell fate is differentially regulated by the microenvironments of the tumor bulk and infiltrative margin. Cell Rep 42, 112472. 10.1016/j.celrep.2023.112472.

4. Xiong, H., Wilson, B.A., Ge, X., Gao, X., Cai, Q., Xu, X., Bachoo, R., and Qin, Z. (2024). Glioblastoma Margin as a Diffusion Barrier Revealed by Photoactivation of Plasmonic Nanovesicles. Nano Letters 24, 1570–1578. 10.1021/acs.nanolett.3c04101.

5. Bastola, S., Pavlyukov, M.S., Yamashita, D., Ghosh, S., Cho, H., Kagaya, N., Zhang, Z., Minata, M., Lee, Y., Sadahiro, H., et al. (2020). Glioma-initiating cells at tumor edge gain signals from tumor core cells to promote their malignancy. Nature Communications 11, 4660. 10.1038/s41467-020-18189-y.

6. Brooks, L.J., and Parrinello, S. (2017). Vascular regulation of glioma stem-like cells: a balancing act. Curr Opin Neurobiol 47, 8–15. 10.1016/j.conb.2017.06.008.

7. Damisah, E.C., Hill, R.A., Rai, A., Chen, F., Rothlin, C.V., Ghosh, S., and Grutzendler, J. (2020). Astrocytes and microglia play orchestrated roles and respect phagocytic territories during neuronal corpse removal in vivo. Sci Adv 6, eaba3239. 10.1126/sciadv.aba3239.

8. Glas, M., Rath, B.H., Simon, M., Reinartz, R., Schramme, A., Trageser, D., Eisenreich, R., Leinhaas, A., Keller, M., Schildhaus, H.U., et al. (2010). Residual tumor cells are unique cellular targets in glioblastoma. Ann Neurol 68, 264–269. 10.1002/ana.22036.

9. Venkataramani, V., Yang, Y., Schubert, M.C., Reyhan, E., Tetzlaff, S.K., Wissmann, N., Botz, M., Soyka, S.J., Beretta, C.A., Pramatarov, R.L., et al. (2022). Glioblastoma hijacks neuronal mechanisms for brain invasion. Cell 185, 2899–2917 e2831. 10.1016/j.cell.2022.06.054.

10. Jayaram, M.A., and Phillips, J.J. (2024). Role of the Microenvironment in Glioma Pathogenesis. Annu Rev Pathol 19, 181–201. 10.1146/annurev-pathmechdis-051122-110348.

11. Yu, K., Hu, Y., Wu, F., Guo, Q., Qian, Z., Hu, W., Chen, J., Wang, K., Fan, X., Wu, X., et al. (2020). Surveying brain tumor heterogeneity by single-cell RNA-sequencing of multi-sector biopsies. Natl Sci Rev 7, 1306–1318. 10.1093/nsr/nwaa099.

12. Hambardzumyan, D., Gutmann, D.H., and Kettenmann, H. (2016). The role of microglia and macrophages in glioma maintenance and progression. Nat Neurosci 19, 20–27. 10.1038/nn.4185.

13. Simmons, G.W., Pong, W.W., Emnett, R.J., White, C.R., Gianino, S.M., Rodriguez, F.J., and Gutmann, D.H. (2011). Neurofibromatosis-1 heterozygosity increases microglia in a spatially and temporally restricted pattern relevant to mouse optic glioma formation and growth. J Neuropathol Exp Neurol 70, 51–62. 10.1097/NEN.0b013e3182032d37.

14. Buonfiglioli, A., and Hambardzumyan, D. (2021). Macrophages and microglia: the cerberus of glioblastoma. Acta Neuropathol Commun 9, 54. 10.1186/s40478-021-01156-z.

15. Gutmann, D.H., and Kettenmann, H. (2019). Microglia/Brain Macrophages as Central Drivers of Brain Tumor Pathobiology. Neuron 104, 442–449. 10.1016/j.neuron.2019.08.028.

16. Pan, Y., and Monje, M. (2022). Neuron–Glial Interactions in Health and Brain Cancer. Advanced Biology 6, 2200122. 10.1002/adbi.202200122.

17. Solomou, G., Young, A.M.H., and Bulstrode, H. (2024). Microglia and macrophages in glioblastoma: landscapes and treatment directions. Mol Oncol. 10.1002/1878-0261.13657.

18. Davalos, D., Grutzendler, J., Yang, G., Kim, J.V., Zuo, Y., Jung, S., Littman, D.R., Dustin, M.L., and Gan, W.B. (2005). ATP mediates rapid microglial response to local brain injury in vivo. Nat Neurosci 8, 752–758. 10.1038/nn1472.

19. Nimmerjahn, A., Kirchhoff, F., and Helmchen, F. (2005). Resting microglial cells are highly dynamic surveillants of brain parenchyma in vivo. Science 308, 1314–1318. 10.1126/science.1110647.

20. Menna, G., Mattogno, P.P., Donzelli, C.M., Lisi, L., Olivi, A., and Della Pepa, G.M. (2022). Glioma-Associated Microglia Characterization in the Glioblastoma Microenvironment through a ’Seed-and Soil’ Approach: A Systematic Review. Brain Sci 12. 10.3390/brainsci12060718.

21. Cuddapah, V.A., Robel, S., Watkins, S., and Sontheimer, H. (2014). A neurocentric perspective on glioma invasion. Nat Rev Neurosci 15, 455–465. 10.1038/nrn3765.

22. Vehlow, A., and Cordes, N. (2013). Invasion as target for therapy of glioblastoma multiforme. Biochim Biophys Acta 1836, 236–244. 10.1016/j.bbcan.2013.07.001.

23. Bayerl, S.H., Niesner, R., Cseresnyes, Z., Radbruch, H., Pohlan, J., Brandenburg, S., Czabanka, M.A., and Vajkoczy, P. (2016). Time lapse in vivo microscopy reveals distinct dynamics of microglia-tumor environment interactions-a new role for the tumor perivascular space as highway for trafficking microglia. Glia 64, 1210–1226. 10.1002/glia.22994.

24. Chen, Z., Ross, J.L., and Hambardzumyan, D. (2019). Intravital 2-photon imaging reveals distinct morphology and infiltrative properties of glioblastoma-associated macrophages. Proc Natl Acad Sci U S A 116, 14254–14259. 10.1073/pnas.1902366116.

25. Resende, F.F., Bai, X., Del Bel, E.A., Kirchhoff, F., Scheller, A., and Titze-de-Almeida, R. (2016). Evaluation of TgH(CX3CR1-EGFP) mice implanted with mCherry-GL261 cells as an in vivo model for morphometrical analysis of glioma-microglia interaction. BMC Cancer 16, 72. 10.1186/s12885-016-2118-3.

26. Johanns, T.M., Ward, J.P., Miller, C.A., Wilson, C., Kobayashi, D.K., Bender, D., Fu, Y., Alexandrov, A., Mardis, E.R., Artyomov, M.N., et al. (2016). Endogenous Neoantigen-Specific CD8 T Cells Identified in Two Glioblastoma Models Using a Cancer Immunogenomics Approach. Cancer Immunol Res 4, 1007–1015. 10.1158/2326-6066.CIR-16-0156.

27. Hicks, W.H., Bird, C.E., Traylor, J.I., Shi, D.D., El Ahmadieh, T.Y., Richardson, T.E., McBrayer, S.K., and Abdullah, K.G. (2021). Contemporary Mouse Models in Glioma Research. Cells 10. 10.3390/cells10030712.

28. Filley, A.C., Henriquez, M., and Dey, M. (2017). Recurrent glioma clinical trial, CheckMate-143: the game is not over yet. Oncotarget 8, 91779–91794. 10.18632/oncotarget.21586.

29. Genoud, V., Marinari, E., Nikolaev, S.I., Castle, J.C., Bukur, V., Dietrich, P.Y., Okada, H., and Walker, P.R. (2018). Responsiveness to anti-PD-1 and anti-CTLA-4 immune checkpoint blockade in SB28 and GL261 mouse glioma models. Oncoimmunology 7, e1501137. 10.1080/2162402X.2018.1501137.

30. Haddad, A.F., Young, J.S., Amara, D., Berger, M.S., Raleigh, D.R., Aghi, M.K., and Butowski, N.A. (2021). Mouse models of glioblastoma for the evaluation of novel therapeutic strategies. Neurooncol Adv 3, vdab100. 10.1093/noajnl/vdab100.

31. Hardcastle, J., Mills, L., Malo, C.S., Jin, F., Kurokawa, C., Geekiyanage, H., Schroeder, M., Sarkaria, J., Johnson, A.J., and Galanis, E. (2017). Immunovirotherapy with measles virus strains in combination with anti-PD-1 antibody blockade enhances antitumor activity in glioblastoma treatment. Neuro Oncol 19, 493–502. 10.1093/neuonc/now179.

32. Wainwright, D.A., Chang, A.L., Dey, M., Balyasnikova, I.V., Kim, C.K., Tobias, A., Cheng, Y., Kim, J.W., Qiao, J., Zhang, L., et al. (2014). Durable therapeutic efficacy utilizing combinatorial blockade against IDO, CTLA-4, and PD-L1 in mice with brain tumors. Clin Cancer Res 20, 5290–5301. 10.1158/1078-0432.CCR-14-0514.

33. Chen, J., Li, Y., Yu, T.S., McKay, R.M., Burns, D.K., Kernie, S.G., and Parada, L.F. (2012). A restricted cell population propagates glioblastoma growth after chemotherapy. Nature 488, 522–526. 10.1038/nature11287.

34. McNicholas, M., De Cola, A., Bashardanesh, Z., Foss, A., Lloyd, C.B., Hébert, S., Faury, D., Andrade, A.F., Jabado, N., Kleinman, C.L., and Pathania, M. (2023). A Compendium of Syngeneic, Transplantable Pediatric High-Grade Glioma Models Reveals Subtype-Specific Therapeutic Vulnerabilities. Cancer Discovery 13, 1592–1615. 10.1158/2159-8290.Cd-23-0004.

35. Pathania, M., De Jay, N., Maestro, N., Harutyunyan, A.S., Nitarska, J., Pahlavan, P., Henderson, S., Mikael, L.G., Richard-Londt, A., Zhang, Y., et al. (2017). H3.3(K27M) Cooperates with Trp53 Loss and PDGFRA Gain in Mouse Embryonic Neural Progenitor Cells to Induce Invasive High-Grade Gliomas. Cancer Cell 32, 684–700 e689. 10.1016/j.ccell.2017.09.014.

36. Zuckermann, M., Hovestadt, V., Knobbe-Thomsen, C.B., Zapatka, M., Northcott, P.A., Schramm, K., Belic, J., Jones, D.T., Tschida, B., Moriarity, B., et al. (2015). Somatic CRISPR/Cas9-mediated tumour suppressor disruption enables versatile brain tumour modelling. Nat Commun 6, 7391. 10.1038/ncomms8391.

37. Brooks, L.J., Clements, M.P., Burden, J.J., Kocher, D., Richards, L., Devesa, S.C., Zakka, L., Woodberry, M., Ellis, M., Jaunmuktane, Z., et al. (2021). The white matter is a pro-differentiative niche for glioblastoma. Nat Commun 12, 2184. 10.1038/s41467-021-22225-w.

38. Claes, A., Idema, A.J., and Wesseling, P. (2007). Diffuse glioma growth: a guerilla war. Acta Neuropathol 114, 443–458. 10.1007/s00401-007-0293-7.

39. Li, Y., Wang, J., Song, S.R., Lv, S.Q., Qin, J.H., and Yu, S.C. (2024). Models for evaluating glioblastoma invasion along white matter tracts. Trends Biotechnol 42, 293–309. 10.1016/j.tibtech.2023.09.005.

40. Scherer, H.J. (1938). Structural Development in Gliomas. The American Journal of Cancer 34, 333–351. 10.1158/ajc.1938.333.

41. Takasaki, K., Abbasi-Asl, R., and Waters, J. (2020). Superficial Bound of the Depth Limit of Two-Photon Imaging in Mouse Brain. eNeuro 7. 10.1523/ENEURO.0255-19.2019.

42. Fuhrmann, F., Nebeling, F.C., Musacchio, F., Mittag, M., Poll, S., Mueller, M., Ambrad Giovannetti, E., Maibach, M., Schaffran, B., Burnside, E., et al. (2024). Three-photon in vivo imaging of neurons and glia in the medial prefrontal cortex with sub-cellular resolution. bioRxiv, 2024.2008.2028.610026. 10.1101/2024.08.28.610026.

43. Horton, N.G., Wang, K., Kobat, D., Clark, C.G., Wise, F.W., Schaffer, C.B., and Xu, C. (2013). In vivo three-photon microscopy of subcortical structures within an intact mouse brain. Nat Photonics 7, 205–209. 10.1038/nphoton.2012.336.

44. Schubert, M.C., Soyka, S.J., Tamimi, A., Maus, E., Schroers, J., Wißmann, N., Reyhan, E., Tetzlaff, S.K., Yang, Y., Denninger, R., et al. (2024). Deep intravital brain tumor imaging enabled by tailored three-photon microscopy and analysis. Nature Communications 15, 7383. 10.1038/s41467-024-51432-4.

45. Neftel, C., Laffy, J., Filbin, M.G., Hara, T., Shore, M.E., Rahme, G.J., Richman, A.R., Silverbush, D., Shaw, M.L., Hebert, C.M., et al. (2019). An Integrative Model of Cellular States, Plasticity, and Genetics for Glioblastoma. Cell 178, 835–849 e821. 10.1016/j.cell.2019.06.024.

46. Bhat, K.P.L., Balasubramaniyan, V., Vaillant, B., Ezhilarasan, R., Hummelink, K., Hollingsworth, F., Wani, K., Heathcock, L., James, J.D., Goodman, L.D., et al. (2013). Mesenchymal differentiation mediated by NF-kappaB promotes radiation resistance in glioblastoma. Cancer Cell 24, 331–346. 10.1016/j.ccr.2013.08.001.

47. Engler, J.R., Robinson, A.E., Smirnov, I., Hodgson, J.G., Berger, M.S., Gupta, N., James, C.D., Molinaro, A., and Phillips, J.J. (2012). Increased microglia/macrophage gene expression in a subset of adult and pediatric astrocytomas. PLoS One 7, e43339. 10.1371/journal.pone.0043339.

48. Kaffes, I., Szulzewsky, F., Chen, Z., Herting, C.J., Gabanic, B., Velázquez Vega, J.E., Shelton, J., Switchenko, J.M., Ross, J.L., McSwain, L.F., et al. (2019). Human Mesenchymal glioblastomas are characterized by an increased immune cell presence compared to Proneural and Classical tumors. OncoImmunology 8, e1655360. 10.1080/2162402X.2019.1655360.

49. Wang, L., Jung, J., Babikir, H., Shamardani, K., Jain, S., Feng, X., Gupta, N., Rosi, S., Chang, S., Raleigh, D., et al. (2022). A single-cell atlas of glioblastoma evolution under therapy reveals cell-intrinsic and cell-extrinsic therapeutic targets. Nat Cancer 3, 1534–1552. 10.1038/s43018-022-00475-x.

50. Aboitiz, F., Scheibel, A.B., Fisher, R.S., and Zaidel, E. (1992). Fiber composition of the human corpus callosum. Brain Res 598, 143–153. 10.1016/0006-8993(92)90178-c.

51. Tong, S., Liu, H., Cheng, H., He, C., Du, Y., Zhuang, Z., Qiu, P., and Wang, K. (2019). Deep-brain three-photon microscopy excited at 1600 nm with silicone oil immersion. Journal of Biophotonics 12, e201800423. 10.1002/jbio.201800423.

52. Venkataramani, V., Schneider, M., Giordano, F.A., Kuner, T., Wick, W., Herrlinger, U., and Winkler, F. (2022). Disconnecting multicellular networks in brain tumours. Nat Rev Cancer 22, 481–491. 10.1038/s41568-022-00475-0.

53. Strojnik, T., Kavalar, R., Zajc, I., Diamandis, E.P., Oikonomopoulou, K., and Lah, T.T. (2009). Prognostic impact of CD68 and kallikrein 6 in human glioma. Anticancer Res 29, 3269–3279.

54. Zhang, J., Li, S., Liu, F., and Yang, K. (2022). Role of CD68 in tumor immunity and prognosis prediction in pan-cancer. Sci Rep 12, 7844. 10.1038/s41598-022-11503-2.

55. Jung, S., Aliberti, J., Graemmel, P., Sunshine, M.J., Kreutzberg, G.W., Sher, A., and Littman, D.R. (2000). Analysis of fractalkine receptor CX(3)CR1 function by targeted deletion and green fluorescent protein reporter gene insertion. Mol Cell Biol 20, 4106–4114. 10.1128/mcb.20.11.4106-4114.2000.

56. Fuhrmann, M., Bittner, T., Jung, C.K., Burgold, S., Page, R.M., Mitteregger, G., Haass, C., LaFerla, F.M., Kretzschmar, H., and Herms, J. (2010). Microglial Cx3cr1 knockout prevents neuron loss in a mouse model of Alzheimer’s disease. Nat Neurosci 13, 411–413. 10.1038/nn.2511.

57. Nebeling, F.C., Poll, S., Justus, L.C., Steffen, J., Keppler, K., Mittag, M., and Fuhrmann, M. (2023). Microglial motility is modulated by neuronal activity and correlates with dendritic spine plasticity in the hippocampus of awake mice. Elife 12. 10.7554/eLife.83176.

58. Banerjee, K., Ratzabi, A., Caspit, I.M., Ganon, O., Blinder, P., Jung, S., and Stein, R. (2023). Distinct spatiotemporal features of microglia and monocyte-derived macrophages in glioma. Eur J Immunol 53, e2250161. 10.1002/eji.202250161.

59. Darmanis, S., Sloan, S.A., Croote, D., Mignardi, M., Chernikova, S., Samghababi, P., Zhang, Y., Neff, N., Kowarsky, M., Caneda, C., et al. (2017). Single-Cell RNA-Seq Analysis of Infiltrating Neoplastic Cells at the Migrating Front of Human Glioblastoma. Cell Rep 21, 1399–1410. 10.1016/j.celrep.2017.10.030.

60. Yabo, Y.A., Moreno-Sanchez, P.M., Pires-Afonso, Y., Kaoma, T., Nosirov, B., Scafidi, A., Ermini, L., Lipsa, A., Oudin, A., Kyriakis, D., et al. (2024). Glioblastoma-instructed microglia transition to heterogeneous phenotypic states with phagocytic and dendritic cell-like features in patient tumors and patient-derived orthotopic xenografts. Genome Med 16, 51. 10.1186/s13073-024-01321-8.

61. Paolicelli, R.C., Sierra, A., Stevens, B., Tremblay, M.E., Aguzzi, A., Ajami, B., Amit, I., Audinat, E., Bechmann, I., Bennett, M., et al. (2022). Microglia states and nomenclature: A field at its crossroads. Neuron 110, 3458–3483. 10.1016/j.neuron.2022.10.020.

62. Stratoulias, V., Venero, J.L., Tremblay, M.E., and Joseph, B. (2019). Microglial subtypes: diversity within the microglial community. EMBO J 38, e101997. 10.15252/embj.2019101997.

63. Bausart, M., Preat, V., and Malfanti, A. (2022). Immunotherapy for glioblastoma: the promise of combination strategies. J Exp Clin Cancer Res 41, 35. 10.1186/s13046-022-02251-2.

64. Yu, M.W., and Quail, D.F. (2021). Immunotherapy for Glioblastoma: Current Progress and Challenges. Front Immunol 12, 676301. 10.3389/fimmu.2021.676301.

65. Osswald, M., Jung, E., Sahm, F., Solecki, G., Venkataramani, V., Blaes, J., Weil, S., Horstmann, H., Wiestler, B., Syed, M., et al. (2015). Brain tumour cells interconnect to a functional and resistant network. Nature 528, 93–98. 10.1038/nature16071.

66. Weil, S., Osswald, M., Solecki, G., Grosch, J., Jung, E., Lemke, D., Ratliff, M., Hanggi, D., Wick, W., and Winkler, F. (2017). Tumor microtubes convey resistance to surgical lesions and chemotherapy in gliomas. Neuro Oncol 19, 1316–1326. 10.1093/neuonc/nox070.

67. Ratliff, M., Karimian-Jazi, K., Hoffmann, D.C., Rauschenbach, L., Simon, M., Hai, L., Mandelbaum, H., Schubert, M.C., Kessler, T., Uhlig, S., et al. (2023). Individual glioblastoma cells harbor both proliferative and invasive capabilities during tumor progression. Neuro Oncol 25, 2150–2162. 10.1093/neuonc/noad109.

68. Wang, X., Liang, J., and Sun, H. (2022). The Network of Tumor Microtubes: An Improperly Reactivated Neural Cell Network With Stemness Feature for Resistance and Recurrence in Gliomas. Front Oncol 12, 921975. 10.3389/fonc.2022.921975.

69. Hausmann, D., Hoffmann, D.C., Venkataramani, V., Jung, E., Horschitz, S., Tetzlaff, S.K., Jabali, A., Hai, L., Kessler, T., Azorin, D.D., et al. (2023). Autonomous rhythmic activity in glioma networks drives brain tumour growth. Nature 613, 179–186. 10.1038/s41586-022-05520-4.

70. Schubert, M.C., Soyka, S.J., Tamimi, A., Maus, E., Denninger, R., Wissmann, N., Reyhan, E., Tetzlaff, S.K., Beretta, C., Drumm, M., et al. (2023). Deep intravital brain tumor imaging enabled by tailored three-photon microscopy and analysis. bioRxiv, 2023.2006.2017.545350. 10.1101/2023.06.17.545350.

71. Weller, J., Potthoff, A.L., Zeyen, T., Schaub, C., Duffy, C., Schneider, M., and Herrlinger, U. (2024). Current status of precision oncology in adult glioblastoma. Mol Oncol. 10.1002/1878-0261.13678.

72. Schneider, M., Vollmer, L., Potthoff, A.L., Ravi, V.M., Evert, B.O., Rahman, M.A., Sarowar, S., Kueckelhaus, J., Will, P., Zurhorst, D., et al. (2021). Meclofenamate causes loss of cellular tethering and decoupling of functional networks in glioblastoma. Neuro Oncol 23, 1885–1897. 10.1093/neuonc/noab092.

73. Belien, A.T., Paganetti, P.A., and Schwab, M.E. (1999). Membrane-type 1 matrix metalloprotease (MT1-MMP) enables invasive migration of glioma cells in central nervous system white matter. J Cell Biol 144, 373–384. 10.1083/jcb.144.2.373.

74. Giger, R.J., Venkatesh, K., Chivatakarn, O., Raiker, S.J., Robak, L., Hofer, T., Lee, H., and Rader, C. (2008). Mechanisms of CNS myelin inhibition: evidence for distinct and neuronal cell type specific receptor systems. Restor Neurol Neurosci 26, 97–115.

75. Gritsenko, P.G., Ilina, O., and Friedl, P. (2012). Interstitial guidance of cancer invasion. J Pathol 226, 185–199. 10.1002/path.3031.

76. Sankowski, R., Böttcher, C., Masuda, T., Geirsdottir, L., Sagar, Sindram, E., Seredenina, T., Muhs, A., Scheiwe, C., Shah, M.J., et al. (2019). Mapping microglia states in the human brain through the integration of high-dimensional techniques. Nature Neuroscience 22, 2098–2110. 10.1038/s41593-019-0532-y.

77. Sankowski, R., Suss, P., Benkendorff, A., Bottcher, C., Fernandez-Zapata, C., Chhatbar, C., Cahueau, J., Monaco, G., Gasull, A.D., Khavaran, A., et al. (2024). Multiomic spatial landscape of innate immune cells at human central nervous system borders. Nat Med 30, 186–198. 10.1038/s41591-023-02673-1.

78. Annovazzi, L., Mellai, M., Bovio, E., Mazzetti, S., Pollo, B., and Schiffer, D. (2018). Microglia immunophenotyping in gliomas. Oncol Lett 15, 998–1006. 10.3892/ol.2017.7386.

79. Chia, K., Mazzolini, J., Mione, M., and Sieger, D. (2018). Tumor initiating cells induce Cxcr4-mediated infiltration of pro-tumoral macrophages into the brain. Elife 7. 10.7554/eLife.31918.

80. Kvisten, M., Mikkelsen, V.E., Stensjoen, A.L., Solheim, O., Van Der Want, J., and Torp, S.H. (2019). Microglia and macrophages in human glioblastomas: A morphological and immunohistochemical study. Mol Clin Oncol 11, 31–36. 10.3892/mco.2019.1856.

81. Mazzolini, J., Le Clerc, S., Morisse, G., Coulonges, C., Zagury, J.F., and Sieger, D. (2022). Wasl is crucial to maintain microglial core activities during glioblastoma initiation stages. Glia 70, 1027–1051. 10.1002/glia.24154.

82. Ricard, C., Tchoghandjian, A., Luche, H., Grenot, P., Figarella-Branger, D., Rougon, G., Malissen, M., and Debarbieux, F. (2016). Phenotypic dynamics of microglial and monocyte-derived cells in glioblastoma-bearing mice. Sci Rep 6, 26381. 10.1038/srep26381.

83. Karabag, D., Scheiblich, H., Griep, A., Santarelli, F., Schwartz, S., Heneka, M.T., and Ising, C. (2023). Characterizing microglial senescence: Tau as a key player. Journal of Neurochemistry 166, 517–533. 10.1111/jnc.15866.

84. Qin, J., Ma, Z., Chen, X., and Shu, S. (2023). Microglia activation in central nervous system disorders: A review of recent mechanistic investigations and development efforts. Front Neurol 14, 1103416. 10.3389/fneur.2023.1103416.

85. Darwish, A., Pammer, M., Gallyas, F., Jr., Vigh, L., Balogi, Z., and Juhasz, K. (2024). Emerging Lipid Targets in Glioblastoma. Cancers (Basel) 16. 10.3390/cancers16020397.

86. Matias, D., Balca-Silva, J., da Graca, G.C., Wanjiru, C.M., Macharia, L.W., Nascimento, C.P., Roque, N.R., Coelho-Aguiar, J.M., Pereira, C.M., Dos Santos, M.F., et al. (2018). Microglia/Astrocytes-Glioblastoma Crosstalk: Crucial Molecular Mechanisms and Microenvironmental Factors. Front Cell Neurosci 12, 235. 10.3389/fncel.2018.00235.

87. van der Poel, M., Ulas, T., Mizee, M.R., Hsiao, C.C., Miedema, S.S.M., Adelia, Schuurman, K.G., Helder, B., Tas, S.W., Schultze, J.L., et al. (2019). Transcriptional profiling of human microglia reveals grey-white matter heterogeneity and multiple sclerosis-associated changes. Nat Commun 10, 1139. 10.1038/s41467-019-08976-7.

88. Grabert, K., Michoel, T., Karavolos, M.H., Clohisey, S., Baillie, J.K., Stevens, M.P., Freeman, T.C., Summers, K.M., and McColl, B.W. (2016). Microglial brain region-dependent diversity and selective regional sensitivities to aging. Nat Neurosci 19, 504–516. 10.1038/nn.4222.

89. Hart, A.D., Wyttenbach, A., Perry, V.H., and Teeling, J.L. (2012). Age related changes in microglial phenotype vary between CNS regions: grey versus white matter differences. Brain Behav Immun 26, 754–765. 10.1016/j.bbi.2011.11.006.

90. Askew, K., Li, K., Olmos-Alonso, A., Garcia-Moreno, F., Liang, Y., Richardson, P., Tipton, T., Chapman, M.A., Riecken, K., Beccari, S., et al. (2017). Coupled Proliferation and Apoptosis Maintain the Rapid Turnover of Microglia in the Adult Brain. Cell Rep 18, 391–405. 10.1016/j.celrep.2016.12.041.

91. Boghozian, R., Sharma, S., Narayana, K., Cheema, M., and Brown, C.E. (2023). Sex and interferon gamma signaling regulate microglia migration in the adult mouse cortex in vivo. Proc Natl Acad Sci U S A 120, e2302892120. 10.1073/pnas.2302892120.

92. Ahn, S.J., Anrather, J., Nishimura, N., and Schaffer, C.B. (2018). Diverse Inflammatory Response After Cerebral Microbleeds Includes Coordinated Microglial Migration and Proliferation. Stroke 49, 1719–1726. 10.1161/STROKEAHA.117.020461.

93. Ren, Y., Huang, Z., Zhou, L., Xiao, P., Song, J., He, P., Xie, C., Zhou, R., Li, M., Dong, X., et al. (2023). Spatial transcriptomics reveals niche-specific enrichment and vulnerabilities of radial glial stem-like cells in malignant gliomas. Nat Commun 14, 1028. 10.1038/s41467-023-36707-6.

94. Dadwal, S., and Heneka, M.T. (2024). Microglia heterogeneity in health and disease. FEBS Open Bio 14, 217–229. 10.1002/2211-5463.13735.

95. Feng, X., Szulzewsky, F., Yerevanian, A., Chen, Z., Heinzmann, D., Rasmussen, R.D., Alvarez-Garcia, V., Kim, Y., Wang, B., Tamagno, I., et al. (2015). Loss of CX3CR1 increases accumulation of inflammatory monocytes and promotes gliomagenesis. Oncotarget 6, 15077–15094. 10.18632/oncotarget.3730.

96. Giordano, F.A., Layer, J.P., Leonardelli, S., Friker, L.L., Turiello, R., Corvino, D., Zeyen, T., Schaub, C., Muller, W., Sperk, E., et al. (2024). L-RNA aptamer-based CXCL12 inhibition combined with radiotherapy in newly-diagnosed glioblastoma: dose escalation of the phase I/II GLORIA trial. Nat Commun 15, 4210. 10.1038/s41467-024-48416-9.

97. Guizar-Sicairos, M., Thurman, S.T., and Fienup, J.R. (2008). Efficient subpixel image registration algorithms. Opt. Lett. 33, 156–158. 10.1364/OL.33.000156.

98. 98. van der Walt, S., Schönberger, J.L., Nunez-Iglesias, J., Boulogne, F., Warner, J.D., Yager, N., Gouillart, E., and Yu, T. (2014). scikit-image: image processing in Python. PeerJ 2, e453. 10.7717/peerj.453.

